# Divergent Pontine and Cerebellar Pathology in Preterm Infants With Necrotizing Enterocolitis

**DOI:** 10.64898/2026.10.30.763262

**Authors:** Georgios Sanidas, Helen Chen, Gabriele Simonti, Javid Ghaemmaghami, Rodolfo Cardenas, Chad Byrd, Courtney Lowe, Laura Kim, Maria Triantafyllou, Nora Wolff, Robinson Vidva, Anthony Sandler, Billie Lou Short, Tania Siahanidou, Christina Kanaka-Gantenbein, Henrik Sahlin Pettersen, Mariam Said, Khodayar Rais-Bahrami, Vittorio Gallo, Frances Northington, Ioannis Koutroulis, Panagiotis Kratimenos

**Author notes:** **Corresponding Author:** Panagiotis Kratimenos, MD, PhD, Children’s National Hospital, George Washington University School of Medicine and Health Sciences, **Email:**.

## Abstract

**Importance:** Preterm survivors of necrotizing enterocolitis (NEC) face substantial neurodevelopmental impairment, but the regional biology and timing of associated brain abnormalities remain poorly defined.

**Objective:** To define the regional, cellular, molecular, and temporal features of NEC-associated brain pathology.

**Design:** Prospective multimodal clinical cohort study (2012–2024), complemented by postmortem tissue analyses.

**Setting:** Level IV neonatal intensive care unit at Children’s National Hospital, Washington, DC.

**Participants:** Of 400 infants screened, 253 preterm infants born before 37 weeks’ gestation (128 with NEC and 125 without NEC) were included across neuroimaging, plasma metabolomic, and postmortem cohorts.

**Exposures:** Medical or surgical NEC (modified Bell stage ≥II).

**Main Outcomes and Measures:** Primary outcomes were term-equivalent magnetic resonance imaging (MRI) abnormalities and serial ultrasound hindbrain growth; secondary outcomes were postmortem histopathology, brain and intestinal transcriptomics, and plasma metabolomics.

**Results:** Among 179 infants with term-equivalent MRI (89 NEC, 90 controls; 102 [57.0%] male; mean [SD] gestational age, 26.3 [2.2] weeks), NEC was associated with a higher prevalence of pontine abnormalities (12.4% vs 1.1%; adjusted prevalence ratio, 9.14; 95% CI, 1.16–71.92), alongside white matter, callosal, and cerebellar abnormalities. In postmortem tissue (n = 27), white matter pathology qualitatively resembled that of prematurity, whereas the hindbrain showed distinct regional phenotypes. The cerebellum exhibited external granular layer contraction (42% reduction; *P* = .04), while the pons showed neuronal injury with increased morphological apoptotic burden (P ≤ .05). Brain and intestinal transcriptomics (n = 14), complemented by plasma metabolomics (n = 29), revealed divergent inflammatory and metabolic signatures. The pons showed increased neuroinflammatory and reduced mitochondrial programs, whereas the cerebellum showed reduced metabolic and cell-cycle programs alongside circulating lysophospholipid depletion. Serial ultrasonography (n = 159), anchored to the cohort mean NEC onset age of 29 weeks’ postmenstrual age, identified earlier cerebellar dimensional deficits (P = .015) and later pontine growth deceleration (*P* < .001).

**Conclusions and Relevance:** In this cohort study, NEC-associated hindbrain pathology showed asynchronous growth patterns accompanied by distinct cellular and molecular phenotypes. These findings support investigation of regional vulnerability and stage-specific neuroprotection; individual injury timing and causal mechanisms remain unresolved.

**Key Points:** *Question:* How does necrotizing enterocolitis (NEC) affect specific brain regions across the clinical course of disease in preterm infants?

*Findings:* In this multimodal cohort study of 253 preterm infants, NEC was associated with asynchronous hindbrain abnormalities. Cerebellar dimensional deficits were detectable before the cohort mean age at NEC onset and persisted without detectable further slowing, alongside progenitor layer contraction and reduced growth-related programs. Pontine growth deceleration emerged after the mean onset age with neuronal injury and inflammatory transcriptomic enrichment.

*Meaning:* Regional differences in growth, cellular pathology, and molecular programs support investigation of stage-specific neuroprotective approaches to NEC-associated brain injury.

## Introduction

Preterm birth forces rapid third-trimester brain development to proceed ex utero, exposing developing neural circuits to the metabolic and inflammatory stress imposed by an immature gastrointestinal tract.^1–4^ This developmental susceptibility is particularly evident in necrotizing enterocolitis (NEC), a life-threatening intestinal disease of prematurity that can burden survivors with long-term neurodevelopmental impairment.^4–7^ Experimental NEC models implicate gut-derived inflammatory signaling in injury to the developing brain, particularly cerebral white matter.^8,9^ Yet clinical NEC does not arise de novo; overt disease can emerge from a prodromal phase marked by gut microbial dysbiosis and intestinal barrier dysfunction.^10–12^ Across this trajectory, evolving disease-related insults may disrupt brain maturation and compromise neuronal integrity, extending the neurological consequences of NEC beyond the acute intestinal episode.^4,10,12^

During the late second and third trimesters, brain regions follow distinct maturational trajectories.^1–3,13^ While glial maturation prepares cerebral white matter for myelination,^1,14,15^ other regions undergo rapid progenitor proliferation and synaptic development.^1–3,16–18^ Experimental evidence links lipid synthesis and nutrient transport to neural progenitor proliferation and brain growth,^19–21^ while NEC models demonstrate susceptibility to inflammatory injury.^8,9^ These processes may shape regional responses to NEC across the disease course. However, integrated human evidence remains scarce, leaving unresolved when NEC-associated brain pathology begins and how disrupted maturation and cellular injury contribute to its evolution.

Here, we examine the regional and temporal organization of NEC-associated brain pathology in a multimodal cohort of 253 preterm infants. We combine systematic whole-brain magnetic resonance imaging (MRI) at term-equivalent age with comparative postmortem neuropathology, multi-omic profiling, and longitudinal cranial ultrasonography to relate regional growth trajectories to cellular and molecular findings. This integration addresses how developmental context and the systemic metabolic environment relate to the timing and nature of NEC-associated brain abnormalities.

## Methods

### Study Design and Cohorts

This multimodal human study integrated a prospective in vivo cohort with a complementary postmortem tissue cohort at Children’s National Hospital between 2012 and 2024, following institutional review board approval and written parental informed consent. Reporting follows the Strengthening the Reporting of Observational Studies in Epidemiology (STROBE) guideline.^22^

Of 400 preterm infants (<37 weeks’ gestation) screened, 147 with congenital malformations, chromosomal abnormalities, or non-NEC intestinal pathology were excluded, yielding 253 unique participants (128 with NEC, 125 without NEC) (eFigure 1). Clinician-recorded sex and parent-reported race and ethnicity (collected to characterize the study population) were extracted from medical records (Table 1; eTable 1). NEC (modified Bell stage ≥II with radiographic confirmation) was stratified into medical (nonoperative) and surgical (laparotomy or peritoneal drainage) management.

**Table 1.**
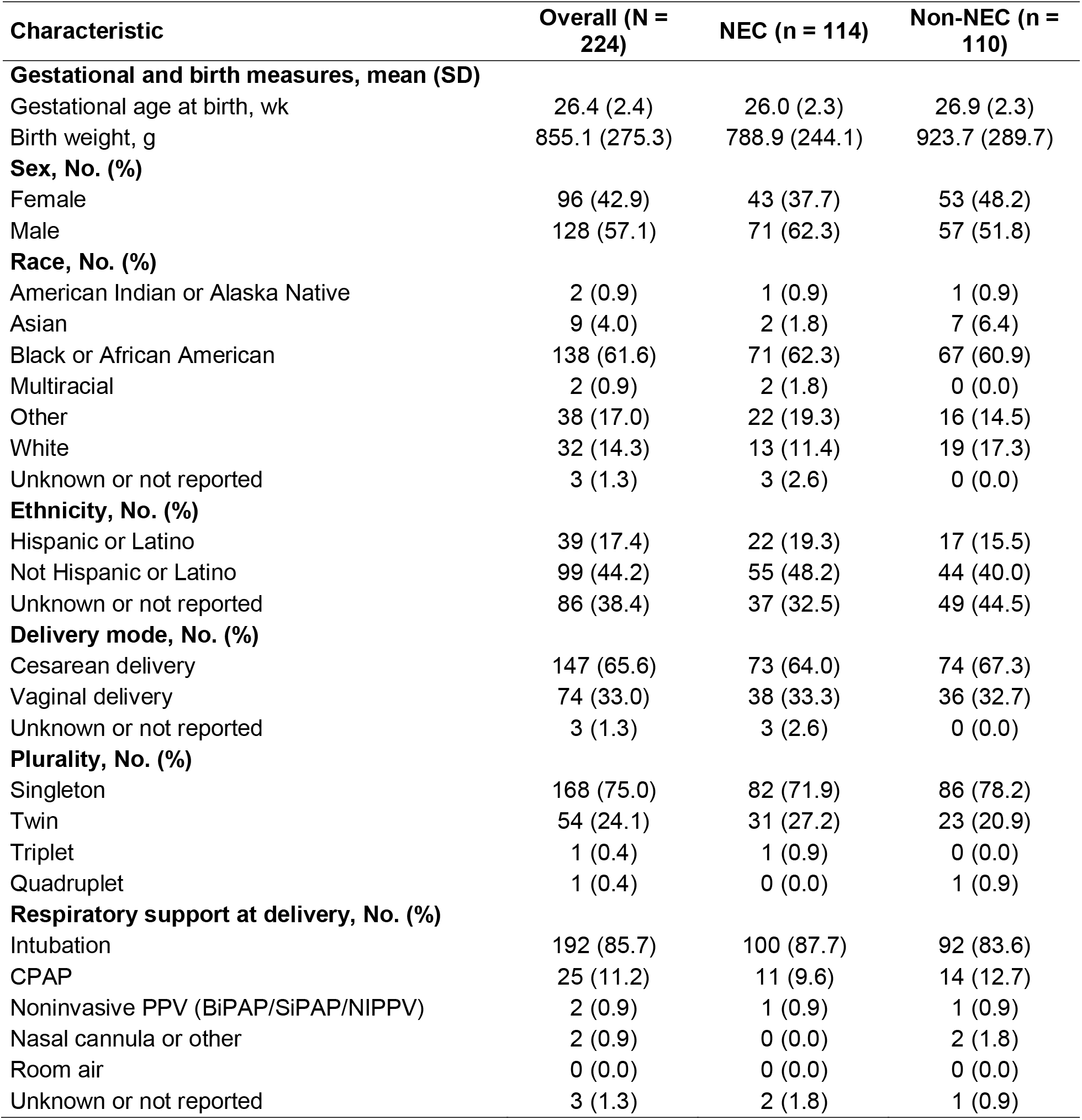
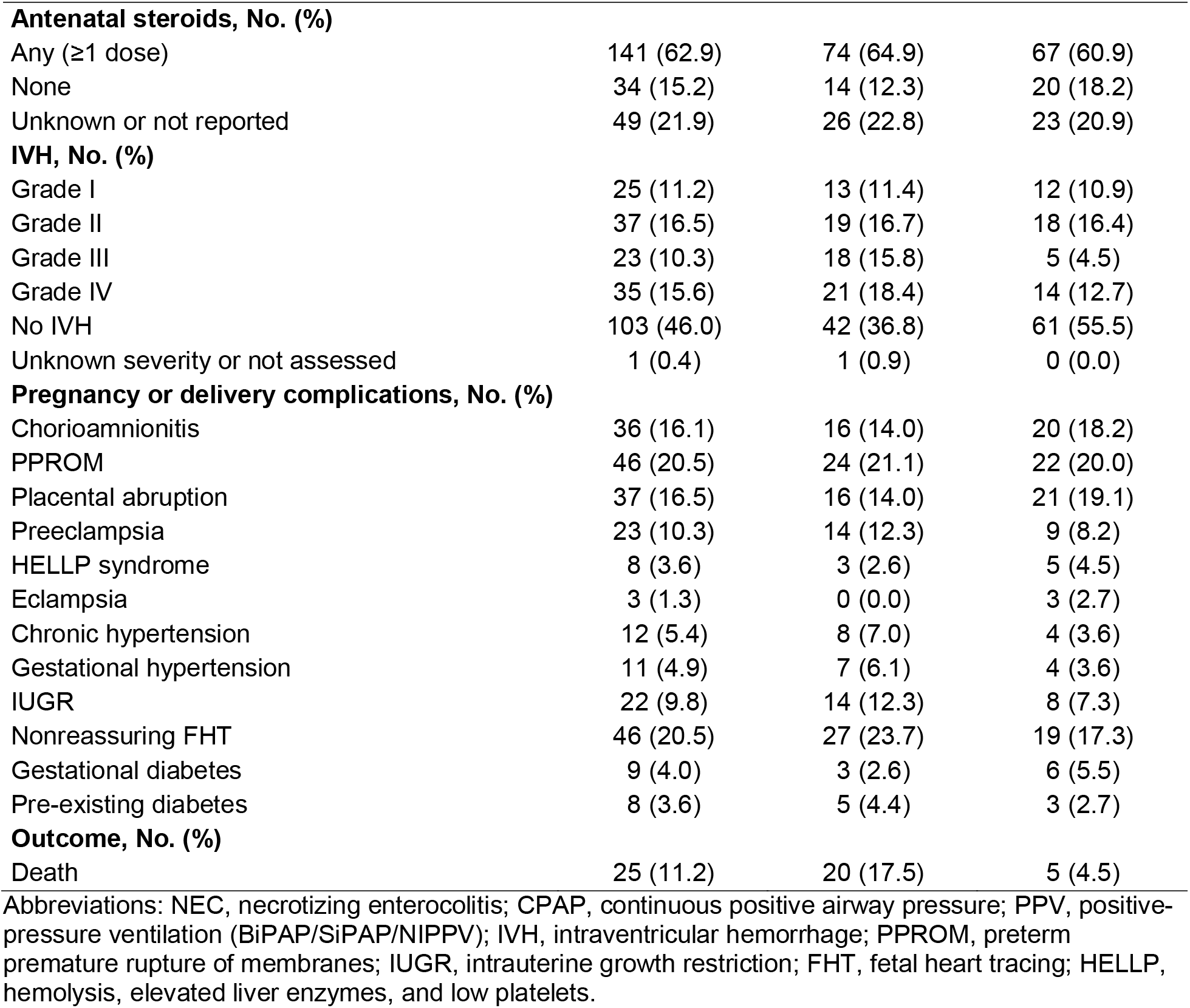
Perinatal and Clinical Characteristics of the In-Vivo Cohort, by NEC Status.

| Characteristic | Overall (N = 224) | NEC (n = 114) | Non-NEC (n = 110) |
|---|---|---|---|
| **Gestational and birth measures, mean (SD)** |  |  |  |
| Gestational age at birth, wk | 26.4 (2.4) | 26.0 (2.3) | 26.9 (2.3) |
| Birth weight, g | 855.1 (275.3) | 788.9 (244.1) | 923.7 (289.7) |
| **Sex, No. (%)** |  |  |  |
| Female | 96 (42.9) | 43 (37.7) | 53 (48.2) |
| Male | 128 (57.1) | 71 (62.3) | 57 (51.8) |
| **Race, No. (%)** |  |  |  |
| American Indian or Alaska Native | 2 (0.9) | 1 (0.9) | 1 (0.9) |
| Asian | 9 (4.0) | 2 (1.8) | 7 (6.4) |
| Black or African American | 138 (61.6) | 71 (62.3) | 67 (60.9) |
| Multiracial | 2 (0.9) | 2 (1.8) | 0 (0.0) |
| Other | 38 (17.0) | 22 (19.3) | 16 (14.5) |
| White | 32 (14.3) | 13 (11.4) | 19 (17.3) |
| Unknown or not reported | 3 (1.3) | 3 (2.6) | 0 (0.0) |
| **Ethnicity, No. (%)** |  |  |  |
| Hispanic or Latino | 39 (17.4) | 22 (19.3) | 17 (15.5) |
| Not Hispanic or Latino | 99 (44.2) | 55 (48.2) | 44 (40.0) |
| Unknown or not reported | 86 (38.4) | 37 (32.5) | 49 (44.5) |
| **Delivery mode, No. (%)** |  |  |  |
| Cesarean delivery | 147 (65.6) | 73 (64.0) | 74 (67.3) |
| Vaginal delivery | 74 (33.0) | 38 (33.3) | 36 (32.7) |
| Unknown or not reported | 3 (1.3) | 3 (2.6) | 0 (0.0) |
| **Plurality, No. (%)** |  |  |  |
| Singleton | 168 (75.0) | 82 (71.9) | 86 (78.2) |
| Twin | 54 (24.1) | 31 (27.2) | 23 (20.9) |
| Triplet | 1 (0.4) | 1 (0.9) | 0 (0.0) |
| Quadruplet | 1 (0.4) | 0 (0.0) | 1 (0.9) |
| **Respiratory support at delivery, No. (%)** |  |  |  |
| Intubation | 192 (85.7) | 100 (87.7) | 92 (83.6) |
| CPAP | 25 (11.2) | 11 (9.6) | 14 (12.7) |
| Noninvasive PPV (BiPAP/SiPAP/NIPPV) | 2 (0.9) | 1 (0.9) | 1 (0.9) |
| Nasal cannula or other | 2 (0.9) | 0 (0.0) | 2 (1.8) |
| Room air | 0 (0.0) | 0 (0.0) | 0 (0.0) |
| Unknown or not reported | 3 (1.3) | 2 (1.8) | 1 (0.9) |
| **Antenatal steroids, No. (%)** |  |  |  |
| Any (≥1 dose) | 141 (62.9) | 74 (64.9) | 67 (60.9) |
| None | 34 (15.2) | 14 (12.3) | 20 (18.2) |
| Unknown or not reported | 49 (21.9) | 26 (22.8) | 23 (20.9) |
| **IVH, No. (%)** |  |  |  |
| Grade I | 25 (11.2) | 13 (11.4) | 12 (10.9) |
| Grade II | 37 (16.5) | 19 (16.7) | 18 (16.4) |
| Grade III | 23 (10.3) | 18 (15.8) | 5 (4.5) |
| Grade IV | 35 (15.6) | 21 (18.4) | 14 (12.7) |
| No IVH | 103 (46.0) | 42 (36.8) | 61 (55.5) |
| Unknown severity or not assessed | 1 (0.4) | 1 (0.9) | 0 (0.0) |
| **Pregnancy or delivery complications, No. (%)** |  |  |  |
| Chorioamnionitis | 36 (16.1) | 16 (14.0) | 20 (18.2) |
| PPROM | 46 (20.5) | 24 (21.1) | 22 (20.0) |
| Placental abruption | 37 (16.5) | 16 (14.0) | 21 (19.1) |
| Preeclampsia | 23 (10.3) | 14 (12.3) | 9 (8.2) |
| HELLP syndrome | 8 (3.6) | 3 (2.6) | 5 (4.5) |
| Eclampsia | 3 (1.3) | 0 (0.0) | 3 (2.7) |
| Chronic hypertension | 12 (5.4) | 8 (7.0) | 4 (3.6) |
| Gestational hypertension | 11 (4.9) | 7 (6.1) | 4 (3.6) |
| IUGR | 22 (9.8) | 14 (12.3) | 8 (7.3) |
| Nonreassuring FHT | 46 (20.5) | 27 (23.7) | 19 (17.3) |
| Gestational diabetes | 9 (4.0) | 3 (2.6) | 6 (5.5) |
| Pre-existing diabetes | 8 (3.6) | 5 (4.4) | 3 (2.7) |
| **Outcome, No. (%)** |  |  |  |
| Death | 25 (11.2) | 20 (17.5) | 5 (4.5) |

Abbreviations: NEC, necrotizing enterocolitis; CPAP, continuous positive airway pressure; PPV, positive-pressure ventilation (BiPAP/SiPAP/NIPPV); IVH, intraventricular hemorrhage; PPROM, preterm premature rupture of membranes; IUGR, intrauterine growth restriction; FHT, fetal heart tracing; HELLP, hemolysis, elevated liver enzymes, and low platelets.

The in vivo cohort (n = 224; 114 NEC, 110 non-NEC) included overlapping subsets for term-equivalent MRI (n = 179), serial cranial ultrasonography (n = 159), and plasma metabolomics (n = 29). The postmortem cohort (n = 32; 17 NEC, 15 non-NEC; including 3 infants from the in vivo cohort who died before discharge) contributed to neuropathology (n = 27) and tissue transcriptomics (n = 14) (eFigure 1; eMethods in the Supplement). Primary outcomes were regional abnormalities on term-equivalent MRI and longitudinal cerebellar and pontine growth trajectories on serial ultrasonography. Secondary outcomes were postmortem cerebellar cortical layer morphometry, pontine cellular density and apoptotic burden, plasma metabolomic profiles at disease onset, and regional tissue transcriptomic pathways.

### Brain MRI Acquisition and Analysis

Assessors blinded to clinical status reviewed 3-T MRI scans obtained at term-equivalent age and coded regional abnormalities as binary indicators across 12 regions; Figure 1 presents 7 selected regions (cortex, white matter, corpus callosum, ventricles, basal ganglia, pons, and cerebellum). Associations with NEC status and surgical vs medical management were evaluated using modified Poisson regression with robust HC0 sandwich variance, adjusted for gestational age at birth and birth weight, and expressed as adjusted prevalence ratios with 95% CIs. Bias-reduced Poisson regression was used for regions with complete separation. Separate lesion-level analyses used unadjusted Firth penalized logistic regression with profile-likelihood 95% CIs (eMethods in the Supplement).

### Cranial Ultrasound Trajectory Modeling

Blinded assessors obtained midsagittal cerebellar and pontine measurements from infants with at least 2 examinations between 22 and 45 weeks’ postmenstrual age (PMA). Regional composite scores averaged component z scores for anteroposterior diameter, height, and cross-sectional area (eMethods in the Supplement). Generalized additive mixed models with infant-level random effects estimated longitudinal trajectories, with statistical separation defined by the pointwise 95% CI for the between-group difference excluding zero. Piecewise mixed-effects models anchored at the cohort mean clinical NEC onset (29 weeks’ PMA) evaluated pre-onset and post-onset slope differences, complemented by linear mixed-effects models testing group-by-age interactions and Welch *t* tests comparing individual growth velocities among infants with at least 3 scans. Biparietal diameter and coronal height growth velocities were compared using analysis of covariance adjusted for midpoint PMA and gestational age at birth.

### Histopathology and Transcriptomics

Formalin-fixed, paraffin-embedded postmortem sections were stained with hematoxylin and eosin (eMethods in the Supplement). Ileal epithelium was segmented using DeepLabV3+ with pathologist verification in QuPath and normalized to muscularis mucosae length. Cerebellar cortical layers (external granular layer, molecular layer, Purkinje cell layer, and internal granular layer) were delineated in QuPath and normalized to molecular layer area divided by thickness. Blinded neuropathologists quantified morphologically intact and karyorrhectic neurons; a morphological apoptotic burden index was calculated for each infant and region as the ratio of karyorrhectic to intact neurons normalized to mean field area (×10^6^). Blinded neuropathologists also performed qualitative histopathological evaluation of the subcortical white matter on H&E-stained sections to assess white matter rarefaction and axonal attenuation. Quantitative cerebellar morphometry included 18 infants (9 NEC, 9 non-NEC; 2-sided exact studentized permutation tests), and pontine morphometry included 14 infants (7 NEC, 7 non-NEC; 2-sided Welch *t* tests), with the infant as the analytic unit.

Bulk RNA sequencing was performed on postmortem ileum (n = 7; 3 NEC, 4 non-NEC), pons (n = 6; 3 NEC, 3 non-NEC), and cerebellum (n = 8; 4 NEC, 4 non-NEC) (eMethods in the Supplement). Reads aligned to GRCh38 were analyzed via DESeq2 Wald tests and multilevel fast gene set enrichment analysis (fgseaMultilevel). Candidate gut-brain ligand-receptor interactions were screened in silico using CellPhoneDB.^23^

### Metabolomics and Cross-Compartment Pathway Integration

Untargeted plasma metabolomics (liquid chromatography–mass spectrometry; Metabolon) included 19 infants with NEC (10 with NEC totalis and 9 with stage III NEC), sampled within 24 hours of clinical onset, and 10 PMA-matched non-NEC controls. Metabolites with more than 50% missing values across the original 30-sample vendor dataset were excluded, retaining 360 metabolites. Vendor-supplied median-normalized, imputed intensities were log₂-transformed and compared in the final 29-cohort using Welch t tests with Benjamini–Hochberg false discovery rate (FDR) correction. Independent plasma metabolomic and tissue transcriptomic datasets were compared across 8 harmonized Reactome, Gene Ontology, and Metabolon pathways using fgsea (eMethods in the Supplement).

### Statistical Analysis

All tests were 2-sided. Nominal *P* < .05 denoted significance for imaging, histopathologic, and cross-compartment pathway comparisons. Differential metabolite abundance required FDR-adjusted *P* < .05 and absolute log_2_ fold change ≥ 1; Hallmark transcriptomic enrichment required FDR-adjusted P < .05. Analyses were conducted in R, version 4.5.3 (R Project for Statistical Computing).

## Results

### Study Population and Clinical Characteristics

The in vivo cohort comprised 224 preterm infants (114 with NEC and 110 without NEC; eFigure 1). Infants with NEC had a lower mean (SD) gestational age at birth (26.0 [2.3] vs 26.9 [2.3] weeks) and birth weight (788.9 [244.1] vs 923.7 [289.7] g). There were 71 male infants (62.3%) in the NEC group and 57 (51.8%) in the non-NEC group. Maternal, obstetric, and additional neonatal characteristics are presented in Table 1.

The postmortem cohort comprised 32 infants (17 with NEC and 15 without NEC). Mean (SD) gestational age and birth weight were 26.4 (3.2) weeks and 890.4 (440.2) g in NEC, compared with 28.5 (4.5) weeks and 1380.4 (1135.5) g in non-NEC infants. The groups included 11 (64.7%) and 6 (40.0%) male infants, respectively (eTable 1).

### Structural Brain Abnormalities and Regional Cellular Pathology

To map the anatomical distribution of NEC-associated brain abnormalities and identify regions of greatest vulnerability, we surveyed term-equivalent MRI in 179 preterm infants, including 89 with NEC (49 medically and 40 surgically managed) and 90 without NEC. Compared with infants without NEC, those with NEC had a higher prevalence of structural abnormalities in the white matter (adjusted prevalence ratio [aPR], 1.55; 95% CI, 1.13–2.12; *P* = .007), corpus callosum (aPR, 2.03; 95% CI, 1.19–3.48; *P* = .010), cerebellum (aPR, 1.69; 95% CI, 1.10–2.61; *P* = .018), and pons (aPR, 9.14; 95% CI, 1.16–71.92; *P* = .036), after adjustment for gestational age at birth and birth weight (Figure 1). Pontine abnormalities showed the largest estimated relative difference between groups, occurring in 11 of 89 infants with NEC (12.4%) compared with 1 of 90 without NEC (1.1%).

**Figure 1.**
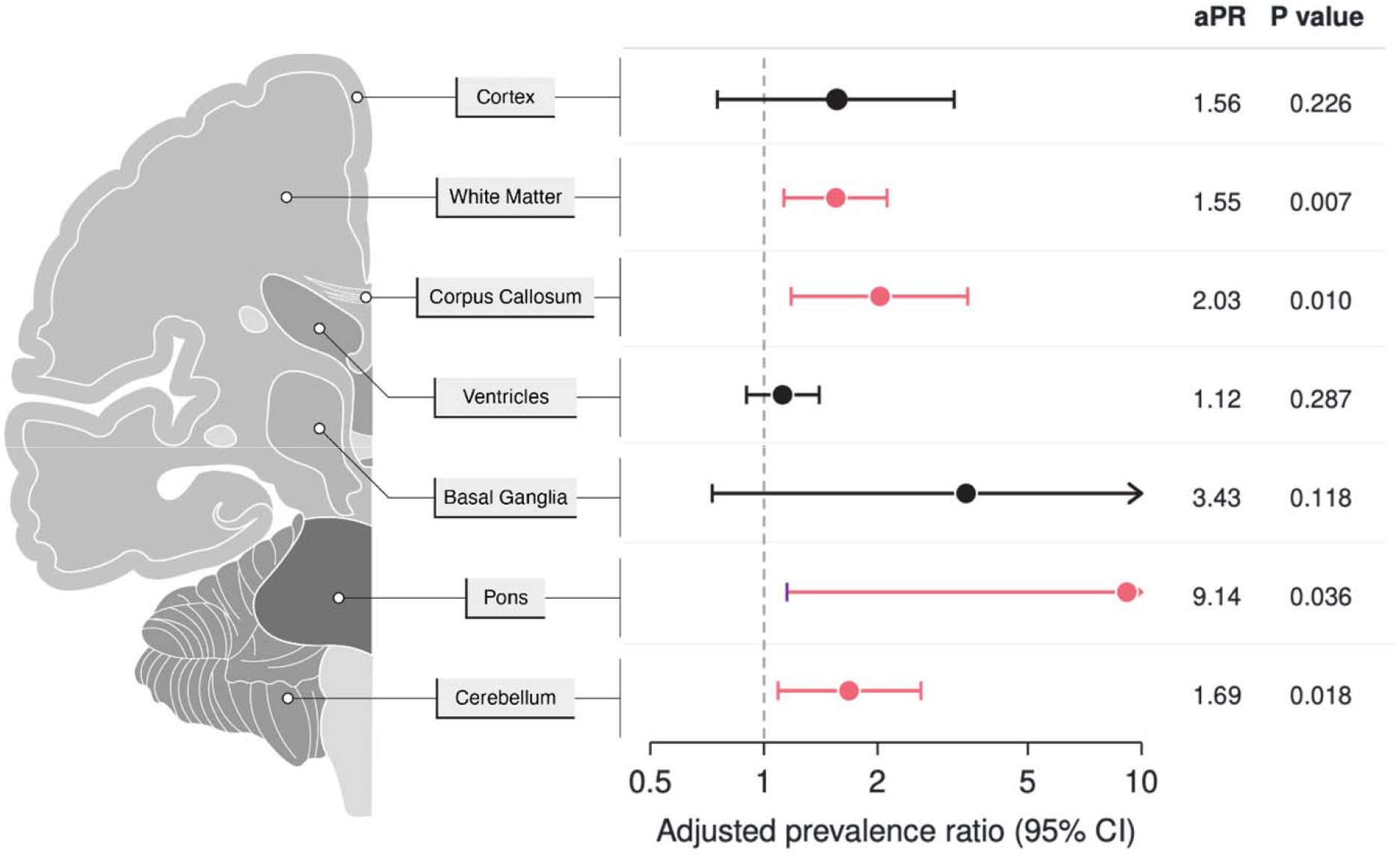
Regional Brain Abnormalities on Term-Equivalent MRI in Infants With Necrotizing Enterocolitis. Anatomical schematic and forest plot showing the prevalence of structural abnormalities across 7 brain regions on term-equivalent MRI in infants with NEC (n = 89) compared with preterm infants without NEC (n = 90). Points represent adjusted prevalence ratios (aPRs), and horizontal lines represent 95% CIs. The vertical dashed line indicates an aPR of 1.0. Red points and lines indicate a statistically significant association (95% CI excludes 1). The rightward-pointing arrow for the pons and basal ganglia indicates that the upper bound of the 95% CI exceeds 10 and has been truncated for visualization. Regional abnormalities were assessed in the cortex, white matter, corpus callosum, ventricles, basal ganglia, pons, and cerebellum. Adjusted prevalence ratios and 95% CIs were estimated using modified Poisson regression with robust HC0 sandwich variance, adjusted for gestational age at birth and birth weight. aPR indicates adjusted prevalence ratio; CI, confidence interval; MRI, magnetic resonance imaging; NEC, necrotizing enterocolitis.

The NEC-associated radiographic pattern included white matter volume loss or hypoplasia and ischemic white matter injury, corpus callosum thinning, and pontine volume loss or hypoplasia (eFigure 2). Within the NEC group, pontine abnormalities occurred in 22.5% of surgically managed infants and 4.1% of medically managed infants (aPR, 5.10; 95% CI, 0.97–26.71; *P* = .05), whereas no difference in cerebellar abnormality prevalence was detected (aPR, 0.92; 95% CI, 0.57–1.49; *P* = .74) (eFigure 3).

To distinguish an amplification of prematurity-associated brain injury from a distinct NEC-associated regional tissue phenotype, we examined postmortem brain and intestinal specimens (eFigure 1). Terminal ileal sections demonstrated mucosal necrosis, pneumatosis intestinalis, and reduced normalized epithelial area in NEC compared with non-NEC controls (*P* = .03) (eFigure 4). Qualitative examination of cerebral subcortical white matter did not identify a distinct NEC-associated lesion pattern (eFigure 5).

In contrast, quantitative hindbrain neuropathology revealed divergent cellular phenotypes in the cerebellum and pons (Figure 2; eFigure 6). In the cerebellar cortex (n = 18), infants with NEC had a 42% smaller normalized external granular layer (EGL) area than non-NEC controls (*P* = .04), with preserved EGL progenitor packing density (*P* = .81), consistent with contraction of the granule progenitor pool (eFigure 6A, 6E). No between-group differences were detected in area or cell density of the internal granular layer (*P* = .10 and *P* = .78), molecular layer (*P* = .06 and *P* = .83), or Purkinje cell layer (*P* = .15 and *P* = .83) (eFigure 6B-D, 6F-H).

**Figure 2.**
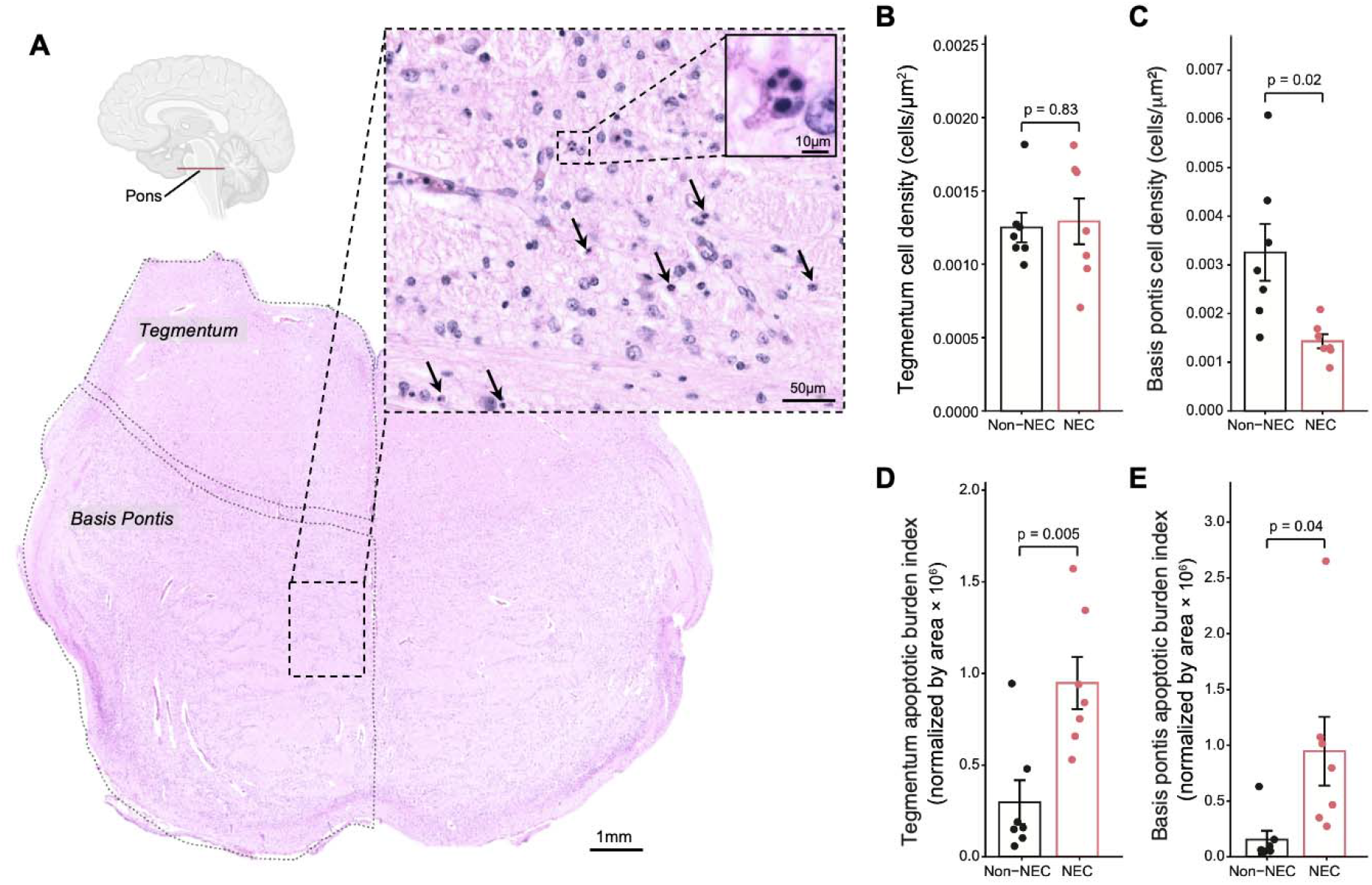
Pontine Cellular Density and Neuronal Injury in Necrotizing Enterocolitis. A, Representative hematoxylin-eosin-stained pontine section delineating the tegmentum and basis pontis, with a sagittal schematic indicating the anatomical plane. Enlarged views demonstrate neuronal somatic shrinkage and karyorrhectic nuclear fragmentation (arrows). Scale bars indicate 1 mm in the overview, 50 μm in the enlarged field, and 10 μm in the inset. B and C, Cellular density in the pontine tegmentum (B; *P* = .83) and basis pontis (C; *P* = .02), expressed as cells per square micrometer. D and E, Morphological apoptotic burden index in the pontine tegmentum (D; *P* = .005) and basis pontis (E; *P* = .04). For each infant and region, the pooled count of karyorrhectic neurons was divided by the pooled count of morphologically normal neurons across evaluable fields; this ratio was normalized to mean field area in square micrometers and multiplied by 10^6^. Quantitative analyses included 7 infants with NEC and 7 without NEC. Each point represents 1 infant; bars indicate group means, and error bars indicate SEM. Black denotes non-NEC infants and red denotes infants with NEC. Between-group comparisons used 2-sided Welch *t* tests. NEC indicates necrotizing enterocolitis.

Among 14 infants undergoing pontine morphometry (7 with NEC and 7 without NEC), NEC specimens showed neuronal injury characterized by eosinophilic somatic shrinkage and karyorrhexis (Figure 2A). While total cellular density in the pontine tegmentum did not differ (*P* = .83; Figure 2B), cellular density in the basis pontis was reduced in NEC (*P* = .02; Figure 2C), accompanied by elevated morphological apoptotic burden index in both the pontine tegmentum (*P* = .005; Figure 2D) and basis pontis (*P* = .04; Figure 2E).

Together, these findings resolve NEC-associated hindbrain pathology into two anatomically adjacent but biologically distinct phenotypes: contraction of the cerebellar progenitor layer and pontine neuronal injury with apoptotic morphology.

### Intestinal–Hindbrain Transcriptional Programs and Systemic Lipid Depletion

To investigate the biological origins of pontine neuronal injury and cerebellar progenitor contraction, we profiled postmortem brain and intestinal transcriptomes to delineate the molecular programs associated with each cellular phenotype.

Pontine transcriptomes showed enrichment of TNF-α/NF-κB signaling (FDR-adjusted *P* < .001; eFigure 7B), paralleling the intestinal inflammatory signature (eFigure 8B). Predicted intestinal–pontine IL6–IL6R and TGFB1–TGFBR1 pairs had more than twofold higher mean interaction-expression scores in NEC (eFigure 9A).

The cerebellum instead showed lower interferon-γ response and apoptosis signatures and reduced programs supporting cell division, cholesterol regulation, and vascular growth (eFigure 10B). Predicted intestinal–cerebellar IGF2 ligand–receptor pairs met the prediction threshold only in non-NEC controls, whereas pairs meeting the threshold only in NEC involved other neurotrophic factors and chemokines (eFigure 9B).

The combination of intestinal and pontine inflammatory enrichment with reduced cerebellar proliferative and lipid-related programs prompted us to examine the circulating metabolic milieu in living infants with NEC and explore its relationship to regional brain responses. Plasma metabolic profiles obtained within 24 hours of NEC diagnosis were most distinct in infants with NEC totalis, although overlap remained among groups (Figure 3A). Among 360 metabolites, 45 differed between groups (33 lower, 12 higher in NEC; FDR-adjusted *P* < .05; absolute log_2_ fold change ≥1; Figure 3B). Lipids accounted for 30 of the 33 metabolites with lower abundance (91%), predominantly lysophosphatidylcholines and lysophosphatidylethanolamines, while cortisol and catecholamine metabolites were elevated (Figure 3B and C).

**Figure 3.**
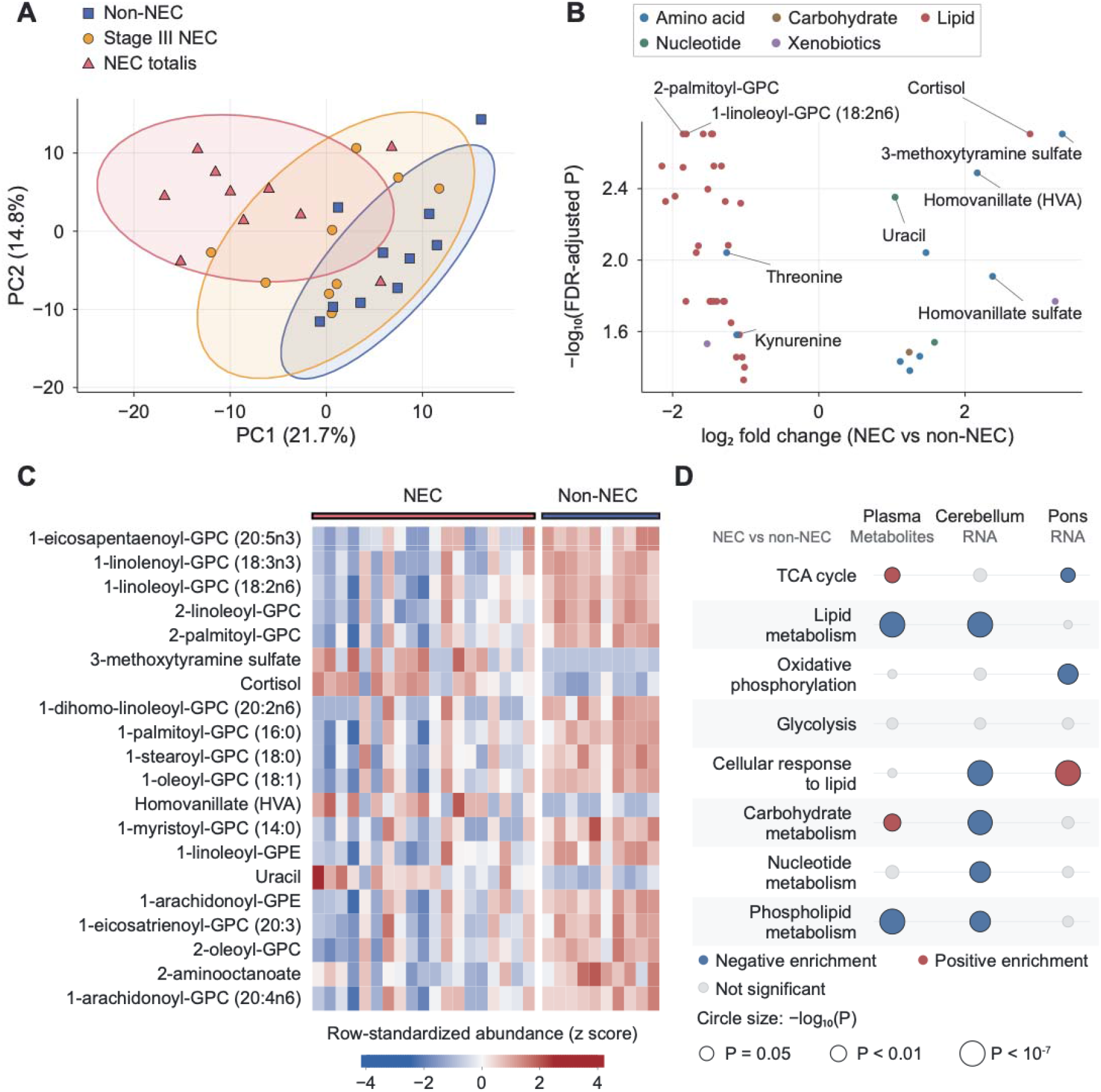
Systemic Lipid Depletion and Region-Specific Molecular Responses in Necrotizing Enterocolitis. A, Principal component analysis of plasma metabolomic profiles from infants with NEC totalis (n = 10), stage III NEC (n = 9), and non-NEC controls (n = 10). Each point represents an infant. NEC samples were obtained within 24 hours of diagnosis; controls were sampled at a matched postmenstrual age. Ellipses show 90% data ellipses. B, Volcano plot of differential plasma metabolite abundance comparing all NEC infants (n = 19) with non-NEC infants (n = 10). Metabolites meeting prespecified significance criteria (FDR-adjusted *P* < .05 and absolute log_2_ fold change ≥ 1) are shown by molecular class. C, Heat map of the 20 significant metabolites with the lowest FDR-adjusted P values. Values represent row-standardized log₂-transformed relative abundance. D, Pathway-level comparison of independently analyzed plasma metabolomic, cerebellar transcriptomic, and pontine transcriptomic profiles. Red and blue indicate positive and negative enrichment, respectively, at nominal P < .05; gray indicates tested nonsignificant pathways. Circle size represents −log₁₀(P), capped at 7. FDR indicates false discovery rate; NEC, necrotizing enterocolitis; NES, normalized enrichment score; PCA, principal component analysis.

Circulating lipid depletion coincided with reduced cerebellar lipid and phospholipid programs, while cellular lipid-response programs were reduced in the cerebellum but increased in the pons (Figure 3D). Carbohydrate- and tricarboxylic acid (TCA) cycle–related metabolite signatures were elevated in plasma, whereas cerebellar carbohydrate and nucleotide programs and pontine mitochondrial programs, including the TCA cycle and oxidative phosphorylation, were reduced (Figure 3D).

Together, these findings identify circulating lysophospholipid depletion alongside intestinal inflammatory enrichment and divergent regional brain responses: inflammatory enrichment and reduced mitochondrial programs in the pons and reduced proliferative and lipid-related programs in the cerebellum.

### Later Severity-Associated Pontine Growth Disruption and Early Cerebellar Dimensional Deficits

To define the timing and evolution of regional growth disturbances during NEC, we next tracked hindbrain growth longitudinally in living infants using serial cranial ultrasonography. In 159 infants (70 with NEC and 89 controls) we examined regional growth relative to the cohort mean NEC onset age of 29 weeks’ postmenstrual age (PMA) (Figure 4A-F).

**Figure 4.**
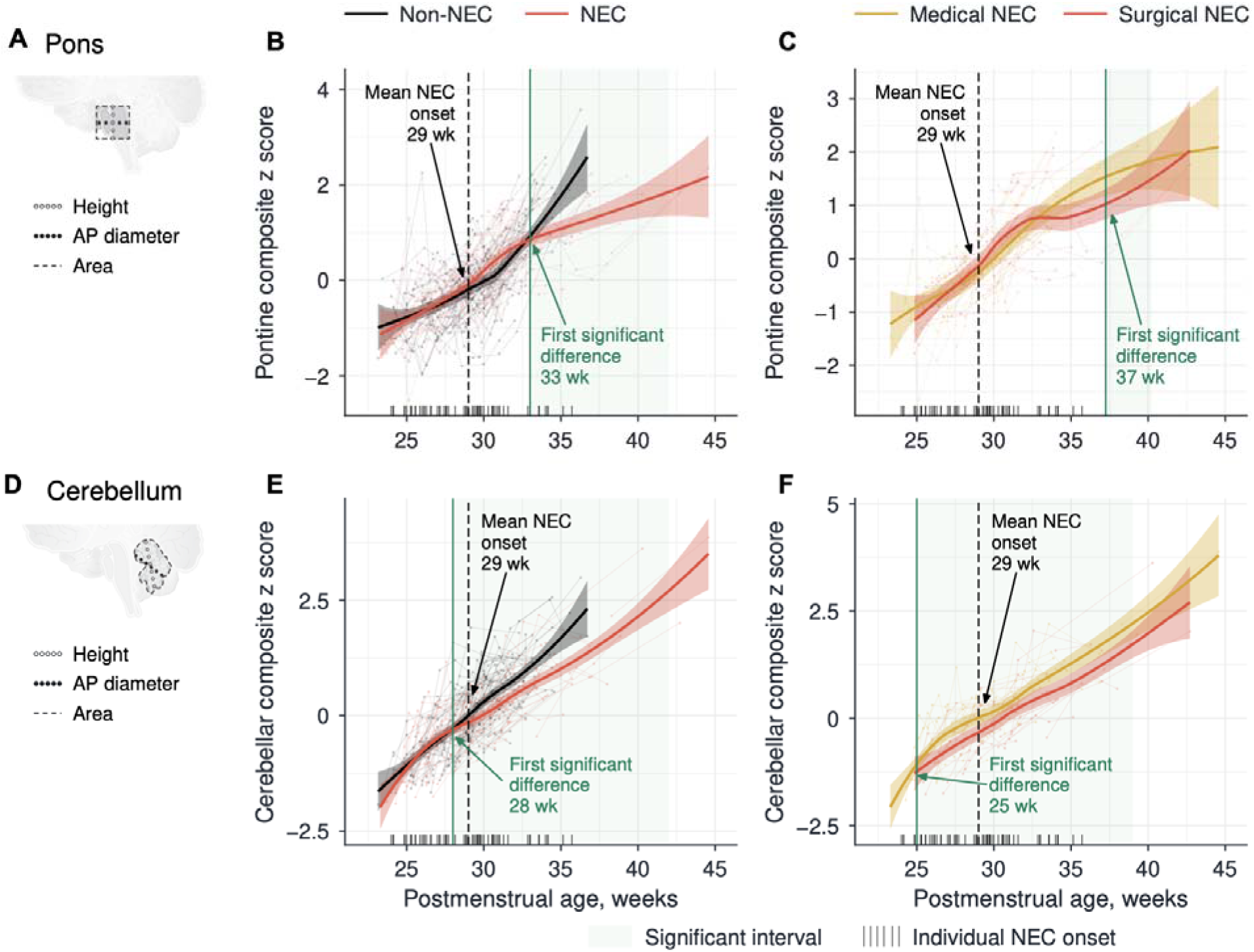
Asynchronous Temporal Trajectories of Pontine and Cerebellar Growth in Necrotizing Enterocolitis. A and D, Midsagittal pontine (A) and cerebellar (D) measurements used to calculate regional composite z scores by averaging standardized height, anteroposterior diameter, and cross-sectional area. B and E, Pontine (B) and cerebellar (E) growth trajectories in 159 infants, comparing 70 infants with necrotizing enterocolitis (NEC; red) with 89 infants without NEC (black). C and F, Pontine (C) and cerebellar (F) growth trajectories among infants with NEC, comparing surgical (n = 34; red) with medical (n = 36; gold) management. Solid curves show descriptive LOESS fits, with surrounding 95% confidence bands. Faint points and connecting lines show individual measurements and within-infant trajectories. Dashed black vertical lines mark the cohort mean NEC onset age of approximately 29 weeks’ postmenstrual age. Rug marks along the horizontal axes indicate individual NEC onset ages. Solid green vertical lines mark the first statistically significant between-group difference. Pale green shading identifies intervals in which the between-group contrast from a separate generalized additive mixed model with infant-level random effects had a pointwise 95% CI excluding zero. The first significant difference indicates statistical detectability and does not establish the onset of the underlying abnormality. AP indicates anteroposterior.

Pontine growth slowing emerged after the cohort mean onset age, with a progressively widening size deficit toward term-equivalent age. Smaller pontine dimensions became detectable at approximately 33 weeks’ PMA and reached a difference of −2.17 z-score units at 40 weeks’ PMA (95% CI, −3.28 to −1.06; *P* < .001) (Figure 4B). Surgically managed infants showed steeper later growth slowing (*P* = .042) and smaller pontine dimensions than medically managed infants from approximately 37 weeks’ PMA (Figure 4C).

Cerebellar dimensions were smaller in infants who developed NEC from approximately 28 weeks’ PMA, before the cohort mean onset age; the mean difference across follow-up was −0.168 z-score units (95% CI, −0.302 to −0.033; *P* = .015) (Figure 4E). Surgically managed infants had smaller cerebellar dimensions than medically managed infants from approximately 25 weeks’ PMA, with a mean difference of −0.323 z-score units (95% CI, −0.495 to −0.152; *P* < .001) (Figure 4F). The surgical–medical cerebellar difference was accompanied by no detectable difference in growth slopes after the cohort mean onset age (P = .793) or in individual growth velocities (P = .846).

Between-group differences in global cranial growth, assessed by biparietal diameter and coronal-height growth rates, did not reach statistical significance (*P* = .778 and *P* = .052, respectively) (eFigure 11).

These trajectories distinguished an early cerebellar dimensional offset from later, progressive pontine growth deceleration that was more pronounced in surgical NEC.

## Discussion

In this multimodal cohort study of 253 preterm infants, we demonstrate that necrotizing enterocolitis (NEC)-associated brain pathology is not a uniform downstream consequence of acute enteric illness, but instead involves asynchronous hindbrain trajectories governed by divergent cellular and molecular programs.

Experimental studies have implicated neuroinflammation in white matter injury,^8,9^ consistent with prominent cerebral white matter abnormalities on clinical MRI in infants with surgical NEC.^24^ The systematic whole-brain imaging survey and regional tissue analyses extend this framework to the developmental organization of the hindbrain. Against a background of white matter changes resembling those of prematurity, the regional findings suggest distinct vulnerabilities of progenitor expansion and neuronal survival.

Serial ultrasonography uncovered a temporal dissociation across the hindbrain, separating an early, persistent cerebellar dimensional deficit from later, progressive pontine growth deceleration. Cerebellar size reductions were detectable before the 29-week cohort mean age at NEC diagnosis, with surgical–medical separation apparent as early as 25 weeks’ PMA. The persistence of this offset without detectable widening raises the possibility that restricted cerebellar expansion and susceptibility to severe intestinal disease share developmental antecedents. In this model, the rapidly proliferating cerebellum may register early inflammatory or metabolic stress that could constrain developmental expansion while also influencing intestinal susceptibility. Disease severity could therefore reflect, in part, the developmental conditions in which NEC emerges. Conversely, pontine growth trajectories diverged later, with progressively widening deficits toward term-equivalent age that were more pronounced in surgical NEC. Whereas the cerebellum may retain the signature of an early constraint on developmental expansion, the pons may reflect vulnerability to the accumulating burden of intestinal injury, systemic inflammation, critical illness, and treatment exposures. These population-level trajectories cannot establish when abnormalities emerged relative to individual NEC onset or whether they contributed to subsequent disease severity.

Lysophospholipid depletion offers a plausible biochemical connection between intestinal inflammation and these regional phenotypes. Phospholipid remodeling couples the maintenance of cellular membranes to innate immune activation and lipid mediator production.^25,26^ Inflammatory changes in lysophosphatidylcholine production and reacylation could therefore contribute to lower circulating abundance.^27^ Dysbiosis and intestinal immune activation can precede clinical recognition of NEC,^11,28^ providing a plausible context for this metabolic disturbance to develop before overt disease. Its onset, and whether it alters lipid availability within the brain, remain unresolved.

The cerebellum places substantial biosynthetic demands on this systemic environment. During late gestation, its growth depends on extensive progenitor expansion and coordinated lineage maturation.^16–18^ Gestational timing organizes these processes,^2^ while prematurity-associated insults can modify their progression.^3^ Experimental evidence links lipid synthesis to adult neural stem and progenitor cell proliferation^19^ and lysophosphatidylcholine-mediated fatty-acid transport to brain growth.^20,21^ The association between early lipid intake and cerebellar growth provides a corresponding clinical link.^29^ Progenitor contraction accompanied by reduced proliferative and lipid-related programs therefore suggests a possible constraint on the metabolic support for tissue expansion.

The pontine findings suggest a different balance between development and injury. Neuron numbers in the pontine nuclei are relatively stable from approximately 27 weeks’ gestation through term,^30^ while pontine growth and circuit maturation continue after birth.^31^ Development in this setting requires preserving established neurons while their connections mature. Inflammatory enrichment and reduced mitochondrial programs accompanying neuronal injury with apoptotic morphology raise the possibility that systemic stress compromises the metabolic support required for neuronal maintenance. Parallel intestinal and pontine inflammatory signatures are consistent with an intestinal contribution to this stress, in keeping with experimental evidence of gut–brain inflammatory signaling.^8,9^ Opposing regional lipid-response programs suggest that the response to metabolic disturbance may deppend on the local tissue context.

Anatomy provides a route through which these distinct cellular vulnerabilities could become a shared functional burden. Pontine nuclei relay cortical input to the cerebellum,^31^ placing pontine neuronal injury within a pathway that supplies the developing cerebellar cortex. Compromised afferent input could therefore compound the consequences of restricted cerebellar expansion. Reported auditory brainstem abnormalities after NEC provide complementary evidence of broader brainstem involvement,^32,33^ although they do not establish pontocerebellar disconnection. Whether impaired cerebellar development and disrupted pontine input interact to influence later disability requires direct investigation.

The translational priority is to resolve the sequence connecting systemic disturbance, regional growth, and functional outcome. Serial ultrasonography provides access to growth over time,^2^ while MRI offers complementary anatomical assessment.^34^ Prospective studies could also test whether cerebellar size measured before clinical diagnosis predicts subsequent NEC or a more severe disease course. Linking these measures to repeated circulating profiles could help distinguish whether inflammatory and lipid changes mark vulnerability, accompany established injury, or reflect its consequences. The molecular findings motivate investigation of lipid homeostasis and inflammatory signaling as complementary targets, while longitudinal validation must establish whether the regional patterns identify distinct and modifiable windows of vulnerability.

### Strengths and Limitations

This study provides a first integrated human account of NEC-associated hindbrain pathology, connecting regional brain tissue findings with longitudinal clinical imaging and intestinal and circulating molecular profiles. These complementary observations establish a foundation for investigating biological processes that cannot be assessed directly and repeatedly in living preterm infants.

Human postmortem tissue provides otherwise inaccessible insight into cellular and molecular pathology, but its availability constrains sample size and each specimen captures a single time point. Plasma profiling likewise reflects the metabolic milieu at diagnosis. Limited overlap between the in vivo and postmortem cohorts prevents reconstruction of a within-infant sequence linking systemic disturbances, tissue pathology, and growth; these observations therefore support biological relationships rather than establish causality. Growth trajectories referenced to the cohort mean NEC onset describe population-level timing, while coexisting neonatal morbidities and intensive care exposures limit attribution of the observed differences to NEC alone.

## Conclusions

NEC-associated hindbrain pathology showed an early cerebellar dimensional offset and later, severity-associated pontine growth deceleration relative to the cohort mean age at NEC onset. These patterns were accompanied by cerebellar progenitor contraction and reduced growth-related programs, and pontine neuronal injury with apoptotic morphology and inflammatory enrichment. The regional distinctions support investigating complementary approaches to preserving maturation and limiting neuronal injury in preterm infants.

## Article Information

### Author Contributions

Dr Kratimenos and Dr Sanidas had full access to all of the data in the study and take responsibility for the integrity of the data and the accuracy of the data analysis. Concept and design: P.K. Acquisition, analysis, or interpretation of data: G.San., H.C., R.C., C.L., G.Sim., L.K., H.S.P., R.V., A.S., J.G., B.L.S., M.S., K.R.-B., M.T., N.W., C.B., P.K. Drafting of the manuscript: G.San., P.K. Critical review of the manuscript for important intellectual content: T.S., C.K.-G., F.N., I.K., V.G., P.K. Statistical analysis: G.San., R.V. Obtained funding: P.K. Administrative, technical, or material support: A.S., B.L.S., M.S., K.R.-B., P.K. Supervision: P.K.

### Data Sharing Statement

Analysis code is available at https://github.com/kratimenoslab/nec-brain-injury. Deidentified processed data supporting the findings are available from the corresponding author upon reasonable request.

### Additional Contributions

We thank the infants included in this study and their families, whose participation made this research possible. We are particularly grateful to the families who consented to postmortem tissue research, contributing to the understanding of brain development and injury in preterm infants.

