## Supplementary material for "Divergent Pontine and Cerebellar Pathology in Preterm Infants With Necrotizing Enterocolitis": Online Supplemental document

Supplementary Online Content

eMethods. Supplementary Methods

eTable 1. Perinatal and Clinical Characteristics of the Postmortem (Autopsy) Cohort, by NEC Status

eFigure 1. Study Cohorts and Analytic Subsets

eFigure 2. Lesion-Level Brain Abnormalities by Anatomical Structure

eFigure 3. Regional Brain Abnormalities in Surgically vs Medically Managed Necrotizing Enterocolitis

eFigure 4. Intestinal Histopathology in the Postmortem Necrotizing Enterocolitis Cohort

eFigure 5. Histopathological Assessment of Cerebral Subcortical White Matter

eFigure 6. Cerebellar Cortical Injury and Morphometry in Necrotizing Enterocolitis

eFigure 7. Pontine Transcriptional Response in Necrotizing Enterocolitis

eFigure 8. Intestinal Transcriptional Programs in Necrotizing Enterocolitis

eFigure 9. Candidate Intestine-to-Hindbrain Ligand-Receptor Signaling in Necrotizing Enterocolitis

eFigure 10. Cerebellar Transcriptional Suppression in Necrotizing Enterocolitis

eFigure 11. Global Cranial Trajectories and Growth Velocities in Necrotizing Enterocolitis

eReferences

### eMethods. Supplementary Methods

#### Study Design and Analytic Cohorts

This multimodal study combined prospective clinical observations and postmortem tissue analyses to identify asynchronous NEC-associated hindbrain abnormalities. Preterm infants with a gestational age of less than 37 weeks were screened. Infants with congenital or chromosomal abnormalities or non-NEC intestinal pathology were excluded. Necrotizing enterocolitis (NEC) was defined as modified Bell stage II or higher with radiographic confirmation. Medical NEC was managed nonoperatively; surgical NEC required laparotomy or peritoneal drainage.

The study included 253 unique infants. The in vivo cohort comprised 224 infants (114 with NEC and 110 without NEC), and the postmortem cohort comprised 32 infants (17 with NEC and 15 without NEC); the cohorts overlapped in 3 participants. Modality-specific analyses used eligible infants with available imaging or specimens. Term-equivalent MRI was available for 179 infants, posterior fossa ultrasound trajectories for 159, and plasma metabolomics for 29. Within the postmortem cohort, 27 infants contributed to neuropathology and 14 to transcriptomics. Tissue availability varied by region, and molecular datasets were not treated as a fully paired, participant-level multi-omics cohort. Cohort flow is presented in eFigure 1. Postmortem cohort characteristics are presented in eTable 1.

No a priori sample-size calculation was performed. Molecular sample sizes were determined by the availability of eligible specimens. Clinical classifications and sample identifiers were reconciled to the final study roster before analysis. MRI abnormalities and longitudinal ultrasound trajectories were the principal imaging outcomes; histopathology and molecular analyses examined the cellular and molecular features of the regional findings.

#### Regional MRI Assessment

##### Image review and outcomes

Term-equivalent 3-T MRI examinations were reviewed by 2 assessors (R.C. and C.L.) blinded to NEC status. Predefined regional abnormalities were coded as present or absent for each infant. Multiple abnormalities in the same region contributed a single positive regional indicator. Figure 1 presents the white matter, corpus callosum, cortex, basal ganglia, ventricles, cerebellum, and pons. The MRI cohort included 89 infants with NEC (49 medical and 40 surgical) and 90 without NEC.

##### Regional prevalence and abnormality types

Regional associations with NEC were estimated using modified Poisson regression with a log link and robust HC0 sandwich variance estimation.^1^ Models included NEC status, gestational age at birth in weeks, and birth weight per 100 g. Exponentiated group coefficients were reported as adjusted prevalence ratios with 95% CIs. Surgical versus medical NEC was evaluated within the NEC subgroup using the same adjustment variables. Bias-reduced Poisson regression was used when complete separation prevented estimation with the standard model.

Additional analyses evaluated clinically grouped abnormality types within regions (eFigure 2). Each abnormality type was represented by a binary indicator, with all infants in the relevant comparison contributing to the denominator. These analyses used unadjusted Firth penalized logistic regression, with odds ratios, profile-likelihood 95% CIs, and penalized likelihood-ratio tests.^2^ The regional surgical–medical comparisons are presented in eFigure 3. Regional prevalence ratios and abnormality-type odds ratios were interpreted as distinct effect measures.

#### Longitudinal Posterior Fossa Ultrasonography

##### Measurements and composite scores

The longitudinal posterior fossa cohort comprised 159 infants: 70 with NEC (36 medical and 34 surgical) and 89 without NEC. Infants contributed at least 2 ultrasound examinations between 22 and 45 weeks’ postmenstrual age (PMA). PMA was calculated as gestational age at birth plus postnatal age in days divided by 7. The dataset contained 546 examinations, with 541 evaluable pontine composite scores and 546 cerebellar composite scores.

Assessors blinded to NEC status and management subgroup obtained measurements using predefined anatomical landmarks and imaging planes. The pontine composite combined the anteroposterior diameter, height, and area of the pons; the cerebellar composite combined the corresponding midsagittal vermian measurements. Each composite was calculated as the mean of its component *z* scores. Measurements of zero or less were treated as missing.

##### Trajectory shape and timing of group differences

Generalized additive mixed models were used to estimate nonlinear relationships between PMA and regional composite scores, accounting for repeated measurements within infants through participant-level random effects. Separate comparisons were performed for NEC versus non-NEC and surgical versus medical NEC. Fitted group-difference curves were used to estimate dimensional contrasts and their 95% CIs across PMA. The earliest evaluated age at which the difference band excluded zero was recorded as the first statistically detectable separation. This threshold describes detectability of a group difference, rather than the biological onset of abnormal growth. Weekly point estimates were accompanied by pointwise 95% CIs.

Clinical NEC onset was defined from the earliest recorded clinical episode classified as medical or surgical NEC. Onset dates were available for 68 of 70 infants with NEC; their mean PMA at onset, 29.0 weeks, provided a common temporal reference. Comparisons with this reference were based on the cohort mean and did not establish that a particular measurement preceded an individual infant’s diagnosis. The pontine contrast at 40 weeks’ PMA was derived from the fitted difference curve. For visualization, Figure 4 displays descriptive LOESS curves and confidence bands; inferential contrasts and intervals of statistical separation were obtained from the generalized additive mixed models.

##### Post-onset slopes, dimensional offsets, and growth velocity

The primary onset-referenced analysis used a piecewise mixed model with a common knot fixed at 29.0 weeks’ PMA. Group contrasts in the post-onset slope component were evaluated separately for the pons and cerebellum and for each NEC comparison. The mean-onset knot was fixed independently of the age at which the fitted group curves first separated statistically.

Complementary linear mixed models distinguished average dimensional differences from changes in those differences over age. Average level differences were estimated from models containing PMA and group as fixed effects, with infant-specific random intercepts and slopes. These models estimated a constant group offset under a common fixed PMA slope. A separate model included PMA centered at 30 weeks, group, and their interaction, again with infant-specific random intercepts and slopes. In this model, the group coefficient estimated the dimensional difference at 30 weeks and the interaction estimated the change in that difference per week.

Individual growth velocities were calculated for infants with at least 3 scans by regressing each infant’s composite score on PMA. The resulting slopes, expressed in *z*-score units per week, were compared between groups using Welch *t* tests.

#### Cranial Growth Measurements

Biparietal diameter and brain coronal height were examined to assess whether the posterior fossa findings were accompanied by differences in more global cranial growth (eFigure 11). These measurements came from a separate 2-examination biometry dataset within the in vivo cohort. Each outcome required a valid pair of measurements and examination ages. After quality review, 62 infants contributed paired biparietal diameter measurements (38 NEC and 24 non-NEC), and 97 contributed paired coronal height measurements (59 NEC and 38 non-NEC). One infant’s paired record was excluded because implausible initial measurements together with unchanged values over a prolonged interval suggested a measurement or date-entry error.

For each infant, growth velocity was calculated as the change in the measurement in millimeters divided by the interval between examinations in weeks. Velocities were compared using analysis of covariance with group, midpoint PMA, and gestational age at birth as predictors. Midpoint PMA was the average PMA of the 2 examinations. Adjusted contrasts were expressed as non-NEC minus NEC in millimeters per week, with 95% CIs. Missing paired measurements were not imputed.

#### Postmortem Histopathology

##### Tissue assessment and deep learning segmentation

Formalin-fixed, paraffin-embedded sections stained with hematoxylin-eosin were examined for intestinal and regional brain pathology. Neuropathologic assessment included 27 infants from the postmortem cohort, with tissue-specific analyses determined by specimen availability. Terminal ileum and cerebellar sections were digitized at ×40 magnification.

Segmentation used DeepLabV3+ convolutional neural networks with an ImageNet-pretrained ResNet-18 encoder (timm, tu-resnet18), output stride 16, atrous spatial pyramid pooling dilation rates of 12, 24, and 36, and batch normalization. Each segmentation task used an ensemble of 3 networks trained iteratively with pathologist correction of model-generated labels in QuPath version 0.5.1.^3^ Models were implemented using PyTorch 2.3.1, segmentation_models_pytorch 0.3.2, timm 0.6.12, and albumentations 2.0.8, with mixed-precision training.

##### Cerebellar segmentation and quantification

Cerebellar cortical layers were manually delineated in QuPath. Analyses included 18 infants, comprising 9 with NEC and 9 without NEC. One infant contributed 2 sections; numeric measurements were averaged within layer across these sections before deriving normalized outcomes. All other infants contributed 1 section.

Raw annotated areas were converted to square micrometers. The normalization factor was calculated as molecular layer area divided by exported molecular layer thickness, yielding a derived length in micrometers. Each layer area was divided by this factor to obtain normalized area, expressed in micrometers.

Between-group comparisons used 2-sided exact studentized permutation tests with the Welch *t* statistic, enumerating all 48,620 allocations of 18 infants into groups of 9. Nominal permutation *P* values are reported in the main text. Mean differences were oriented as NEC minus non-NEC, and reported 95% CIs were calculated using the Welch method. Figures display individual infants and group means with SEM.

Sensitivity analyses applied Benjamini–Hochberg correction across the 4 normalized layer-area comparisons and linear models with HC3 robust standard errors adjusting for developmental age, with additional adjustment for sex. The EGL comparison had an FDR-adjusted P value of .114. Estimates remained negative after adjustment for PMA at death alone (P = .067) and with sex (P = .062), although confidence intervals included zero.

##### Intestinal segmentation and quantification

A first model segmented the intestinal wall into lumen/background, mucosa-submucosa, and muscularis mucosae. A second model segmented epithelium versus nonepithelium and was initialized from a model trained on the public IBDColEpi dataset (doi:10.18710/TLA01U). Both tasks used 1024-pixel patches at 4-fold downsampling. Epithelial area was normalized to muscularis mucosae length. Quantitative intestinal analyses included 11 cases, comprising 5 with NEC and 6 without NEC. For each case, epithelial areas were summed across available images and divided by the summed muscularis mucosae lengths, yielding one area-to-length ratio per case. Images from the same case were treated as technical subsamples. A zero epithelial-area value recorded in the sample key for one NEC case was retained in the analysis. Between-group differences were evaluated using an exact 2-sided studentized permutation test, with the difference in group means divided by its unequal-variance standard error as the test statistic. All 462 possible allocations preserving the group sizes were enumerated. The *P* value was calculated as the proportion of allocations yielding an absolute test statistic at least as large as that observed. Graphs display individual case values and group means with SEM.

##### Pontine neuronal density and apoptotic burden

Quantitative pontine analyses included 14 infants, with 7 in each group. The basis pontis and pontine tegmentum were evaluated separately. Cellular density was quantified using StarDist-based nuclear detection^4^ and expressed as cells per square micrometer, with one summarized value per infant and region. For assessment of apoptotic burden, neurons were distinguished from nonneuronal cells by morphology. Karyorrhectic and morphologically normal neurons were counted manually in 4 fields per infant and region, with morphological review by a neuropathologist blinded to NEC status. Counts were pooled across evaluable fields, and the ratio of karyorrhectic to morphologically normal neurons was divided by the mean evaluable field area in square micrometers and multiplied by 10⁶. This measure represents an area-normalized burden index, rather than the percentage of neurons undergoing apoptosis. One field with an implausible recorded area was excluded following data review; the affected infant remained in the analysis using the other 3 fields. The infant was the unit of analysis. Separate 2-sided Welch *t* tests compared cellular density and apoptotic burden between NEC and non-NEC groups in each pontine region. Individual fields were treated as technical subsamples. The 4 pontine comparisons used nominal *P* values without multiplicity adjustment.

#### Tissue Transcriptomics

##### Tissue subsets and differential expression

Bulk RNA sequencing included intestinal tissue from 7 infants (3 NEC and 4 non-NEC), pontine tissue from 6 (3 NEC and 3 non-NEC), and cerebellar tissue from 8 (4 NEC and 4 non-NEC). These region-specific subsets represented 14 unique infants, with partial overlap across tissues. Reads were aligned to the human GRCh38 reference genome. Analyses compared NEC with non-NEC separately within each tissue.

Gene-level count data were analyzed with DESeq2 using Wald tests for the NEC–non-NEC contrast.^5^ Gene-level *P* values were adjusted using the Benjamini–Hochberg procedure within each tissue.^6^ Differential expression was evaluated at a false discovery rate (FDR) below .05. For pathway analyses, genes were ranked by their DESeq2 Wald statistic, preserving both the direction and strength of the NEC association.

##### Pathway enrichment and expression displays

Preranked gene-set enrichment analysis was performed with fgsea.^7^ Tissue-specific Hallmark analyses evaluated the 50 Hallmark gene sets, with FDR-adjusted *P* values below .05 defining significant enrichment. Positive normalized enrichment scores indicated enrichment toward genes increased in NEC, whereas negative scores indicated enrichment toward genes decreased in NEC. Hallmark displays in eFigures 7, 8, and 10 used the original tissue-specific enrichment outputs. The cross-platform pathway comparison in Figure 3D used gene rankings recomputed from the raw counts, as described below.

Expression heatmaps used transcripts per million transformed as log₂(TPM + 1) and standardized within each gene across samples. The pontine heatmap included the 6 pontine specimens and leading-edge genes from the TNFα signaling via NF-κB Hallmark set. Cerebellar and intestinal heatmaps displayed the corresponding tissue-specific gene panels.

#### Plasma Metabolomics

##### Sampling and clinical groups

The final plasma cohort comprised 29 infants: 10 with NEC totalis, 9 with stage III NEC, and 10 non-NEC controls. NEC samples were collected within 24 hours of clinical onset. Controls were preterm infants who did not develop NEC, with plasma collection timed to match the PMA at NEC onset in cases. Comparisons between NEC and non-NEC groups were unpaired.

##### Profiling and preprocessing

Untargeted plasma metabolomic profiling was performed by Metabolon using liquid chromatography–mass spectrometry. The vendor dataset contained 390 named metabolites spanning amino acid, peptide, carbohydrate, energy, lipid, nucleotide, cofactor, and xenobiotic pathways. Raw ion counts for each metabolite were rescaled to a median of 1.0. Scaled intensities were summarized by clinical group using means, medians, IQRs, and ranges. Intensities represented relative abundance rather than absolute plasma concentrations.

##### Metabolites with more than 50% missing values across the original 30-sample vendor dataset were excluded, retaining 360 of 390 metabolites. The vendor-supplied median-normalized, imputed intensity matrix was log₂-transformed. Differential abundance was evaluated in the final 29-infant cohort after reconciliation of eligibility and NEC versus non-NEC status with the clinical roster.

##### Differential abundance and visualization

NEC and non-NEC log₂ intensities were compared using 2-sided Welch *t* tests, with Benjamini–Hochberg adjustment across the 360 tested metabolites.^6^ Differential abundance required both an adjusted *P* value below .05 and an absolute log₂ fold change of at least 1. Fold-change direction was defined as NEC relative to non-NEC. Principal component analysis was performed on centered and scaled log₂ intensities to visualize overall profile variation. The heatmap displayed the 20 significant metabolites with the lowest adjusted *P* values. Metabolite classes were assigned using the vendor’s superpathway and subpathway annotations.

#### Cross-Platform Pathway Analysis

Plasma metabolomic and regional brain transcriptomic datasets were analyzed independently and compared at the pathway level (Figure 3D). Separate fgsea analyses ranked metabolites by the NEC–non-NEC log₂ fold change and pontine and cerebellar genes by the corresponding DESeq2 Wald statistic. Metabolite sets were defined using Metabolon annotations; transcript sets were obtained from Reactome and Gene Ontology Biological Process collections through msigdbr. Eight pathway categories were harmonized as follows:

TCA cycle. Metabolon “TCA Cycle” was mapped to Reactome “Citric Acid Cycle (TCA Cycle).”

Lipid metabolism. The Metabolon “Lysolipid” subpathway was mapped to Reactome “Metabolism of Lipids”; the plasma set therefore represented lysolipids within the broader transcriptional lipid category.

Oxidative phosphorylation. The Metabolon “Oxidative Phosphorylation” annotation was mapped to Gene Ontology “Oxidative Phosphorylation,” where evaluable.

Glycolysis. Metabolon “Glycolysis, Gluconeogenesis, and Pyruvate Metabolism” was mapped to Reactome “Glucose Metabolism.”

Cellular response to lipid. Gene Ontology “Cellular Response to Lipid” was evaluated in the transcriptomic datasets only.

Carbohydrate metabolism. The Metabolon “Carbohydrate” superpathway was mapped to Gene Ontology “Carbohydrate Metabolic Process.”

Nucleotide metabolism. The Metabolon “Nucleotide” superpathway was mapped to Reactome “Metabolism of Nucleotides.”

Phospholipid metabolism. Metabolon “Phospholipid Metabolism” and “Lysolipid” subpathways were combined and mapped to Reactome “Phospholipid Metabolism.”

The upper gene-set size limit was 1000, allowing inclusion of the specified broad lipid-related sets. Enrichment direction and normalized enrichment scores were compared across platforms. This exploratory 8-pathway analysis used nominal *P* values, with *P* < .05 indicating enrichment in the displayed comparison; these *P* values were distinct from the FDR-adjusted Hallmark results. Pathways without an evaluable set were not assigned an enrichment estimate. Cross-platform concordance was assessed at the level of biological programs, without fitting a participant-level joint model.

#### Cross-Tissue Ligand–Receptor Analysis

Cross-tissue ligand–receptor relationships were evaluated using CellPhoneDB and its curated interaction and multimeric-complex annotations.^8^ Bulk intestinal, pontine, and cerebellar transcriptomic profiles were analyzed separately for intestine–pons and intestine–cerebellum relationships within NEC and non-NEC conditions. Candidate pairs were assessed using condition-specific permutation testing. For the intestine–pons analysis, pairs significant in both conditions were further prioritized by greater than 2-fold differences in expression, alongside selected interactions from functional-category summaries. For the intestine–cerebellum analysis, candidate pairs were classified according to condition-specific statistical support.

Selected relationships were visualized using chord diagrams (eFigure 9), with expression contrasts and condition-specific classifications used to describe their patterns across NEC and non-NEC tissue. Permutation *P* values represented statistical support within each condition; between-condition expression contrasts were reported separately. Full CellPhoneDB complex identities were retained when interpreting abbreviated figure labels.

#### Statistical Reporting and Software

All statistical tests were 2-sided. Imaging and pontine histopathologic comparisons used nominal *P* values, with *P* < .05 considered statistically significant. Benjamini–Hochberg adjustment was applied separately to differential expression within each tissue, Hallmark enrichment within each tissue, and the 360 metabolite comparisons. The harmonized 8-pathway comparison used nominal *P* values. Estimates were reported with 95% CIs where available. For posterior fossa outcomes, contrasts were oriented as NEC minus non-NEC or surgical minus medical NEC; for cranial growth velocities, the reported contrast was non-NEC minus NEC.

Analyses used the observations available for each outcome, with modality-specific denominators. Serial measurements and histologic fields were handled within infant, rather than counted as additional participants. Metabolomic missing-value imputation used the vendor-supplied matrix. Missing ultrasound measurements were not replaced with zero, and incomplete paired cranial measurements did not contribute to the corresponding velocity analysis.

Analyses were performed in R, version 4.5.3. The analysis environment included brglm2, version 1.1.0; logistf, version 1.26.1; mgcv, version 1.9-4; nlme, version 3.1-168; sandwich, version 3.1-1; DESeq2, version 1.50.2; fgsea, version 1.36.2; and msigdbr, version 26.1.0.

### eTable 1. Perinatal and Clinical Characteristics of the Postmortem (Autopsy) Cohort, by NEC Status

| **Characteristic** | **Overall (N=32)** | **NEC (n=17)** | **Non-NEC (n=15)** |
| --- | --- | --- | --- |
| **Gestational and birth measures, mean (SD)** | | | |
| Gestational age at birth, wk | 27.4 (4.0) | 26.4 (3.2) | 28.5 (4.5) |
| Birth weight, g | 1120.1 (862.6) | 890.4 (440.2) | 1380.4 (1135.5) |
| **Age at death** | | | |
| Postmenstrual age at death, wk, mean (SD) | 33 (10.7) | 35.9 (13.7) | 29.6 (4.2) |
| Postmenstrual age at death, wk, median (range) | 29.9 (23.4–76.0) | 31.4 (23.4–76.0) | 28.7 (24.5–37.0) |
| **Sex, No. (%)** | | | |
| Female | 15 (46.9) | 6 (35.3) | 9 (60.0) |
| Male | 17 (53.1) | 11 (64.7) | 6 (40.0) |
| **Race, No. (%)** | | | |
| American Indian or Alaska Native | 0 (0.0) | 0 (0.0) | 0 (0.0) |
| Asian | 1 (3.1) | 0 (0.0) | 1 (6.7) |
| Black or African American | 18 (56.3) | 10 (58.8) | 8 (53.3) |
| Multiracial | 2 (6.3) | 2 (11.8) | 0 (0.0) |
| Other | 4 (12.5) | 2 (11.8) | 2 (13.3) |
| White | 5 (15.6) | 1 (5.9) | 4 (26.7) |
| Unknown or not reported | 2 (6.3) | 2 (11.8) | 0 (0.0) |
| **Ethnicity, No. (%)** | | | |
| Hispanic or Latino | 5 (15.6) | 2 (11.8) | 3 (20.0) |
| Not Hispanic or Latino | 4 (12.5) | 3 (17.6) | 1 (6.7) |
| Unknown or not reported | 23 (71.9) | 12 (70.6) | 11 (73.3) |
| **Delivery mode, No. (%)** | | | |
| Cesarean delivery | 22 (68.8) | 11 (64.7) | 11 (73.3) |
| Vaginal delivery | 9 (28.1) | 5 (29.4) | 4 (26.7) |
| Unknown or not reported | 1 (3.1) | 1 (5.9) | 0 (0.0) |
| **Plurality, No. (%)** | | | |
| Singleton | 27 (84.4) | 14 (82.4) | 13 (86.7) |
| Twin | 5 (15.6) | 3 (17.6) | 2 (13.3) |
| Triplet | 0 (0.0) | 0 (0.0) | 0 (0.0) |
| Quadruplet | 0 (0.0) | 0 (0.0) | 0 (0.0) |
| **Respiratory support at delivery, No. (%)** | | | |
| Intubation | 31 (96.9) | 17 (100.0) | 14 (93.3) |
| CPAP | 1 (3.1) | 0 (0.0) | 1 (6.7) |
| Noninvasive PPV (BiPAP/SiPAP/NIPPV) | 0 (0.0) | 0 (0.0) | 0 (0.0) |
| Nasal cannula or other | 0 (0.0) | 0 (0.0) | 0 (0.0) |
| Room air | 0 (0.0) | 0 (0.0) | 0 (0.0) |
| Unknown or not reported | 0 (0.0) | 0 (0.0) | 0 (0.0) |
| **Antenatal steroids, No. (%)** | | | |
| Any (≥1 dose) | 19 (59.4) | 10 (58.8) | 9 (60.0) |
| None | 5 (15.6) | 3 (17.6) | 2 (13.3) |
| Unknown or not reported | 8 (25.0) | 4 (23.5) | 4 (26.7) |
| **IVH, No. (%)** | | | |
| Grade I | 1 (3.1) | 0 (0.0) | 1 (6.7) |
| Grade II | 5 (15.6) | 4 (23.5) | 1 (6.7) |
| Grade III | 1 (3.1) | 1 (5.9) | 0 (0.0) |
| Grade IV | 4 (12.5) | 0 (0.0) | 4 (26.7) |
| No IVH | 18 (56.3) | 11 (64.7) | 7 (46.7) |
| Unknown severity or not assessed | 3 (9.4) | 1 (5.9) | 2 (13.3) |
| Severe IVH (grade III-IV) | 5 (15.6) | 1 (5.9) | 4 (26.7) |
| **Pregnancy or delivery complications, No. (%)** | | | |
| Chorioamnionitis | 5 (15.6) | 1 (5.9) | 4 (26.7) |
| PPROM | 10 (31.3) | 5 (29.4) | 5 (33.3) |
| Placental abruption | 3 (9.4) | 1 (5.9) | 2 (13.3) |
| Preeclampsia | 6 (18.8) | 4 (23.5) | 2 (13.3) |
| HELLP syndrome | 0 (0.0) | 0 (0.0) | 0 (0.0) |
| Eclampsia | 0 (0.0) | 0 (0.0) | 0 (0.0) |
| Chronic hypertension | 0 (0.0) | 0 (0.0) | 0 (0.0) |
| Gestational hypertension | 2 (6.3) | 1 (5.9) | 1 (6.7) |
| IUGR | 3 (9.4) | 1 (5.9) | 2 (13.3) |
| Nonreassuring FHT | 10 (31.3) | 3 (17.6) | 7 (46.7) |
| Gestational diabetes | 2 (6.3) | 1 (5.9) | 1 (6.7) |
| Pre-existing diabetes | 1 (3.1) | 1 (5.9) | 0 (0.0) |

Abbreviations: NEC, necrotizing enterocolitis; CPAP, continuous positive airway pressure; PPV, positive-pressure ventilation (BiPAP/SiPAP/NIPPV); IVH, intraventricular hemorrhage; PMA, postmenstrual age; PPROM, preterm premature rupture of membranes; IUGR, intrauterine growth restriction; FHT, fetal heart tracing; HELLP, hemolysis, elevated liver enzymes, and low platelets.

### eFigure 1. Study Cohorts and Analytic Subsets

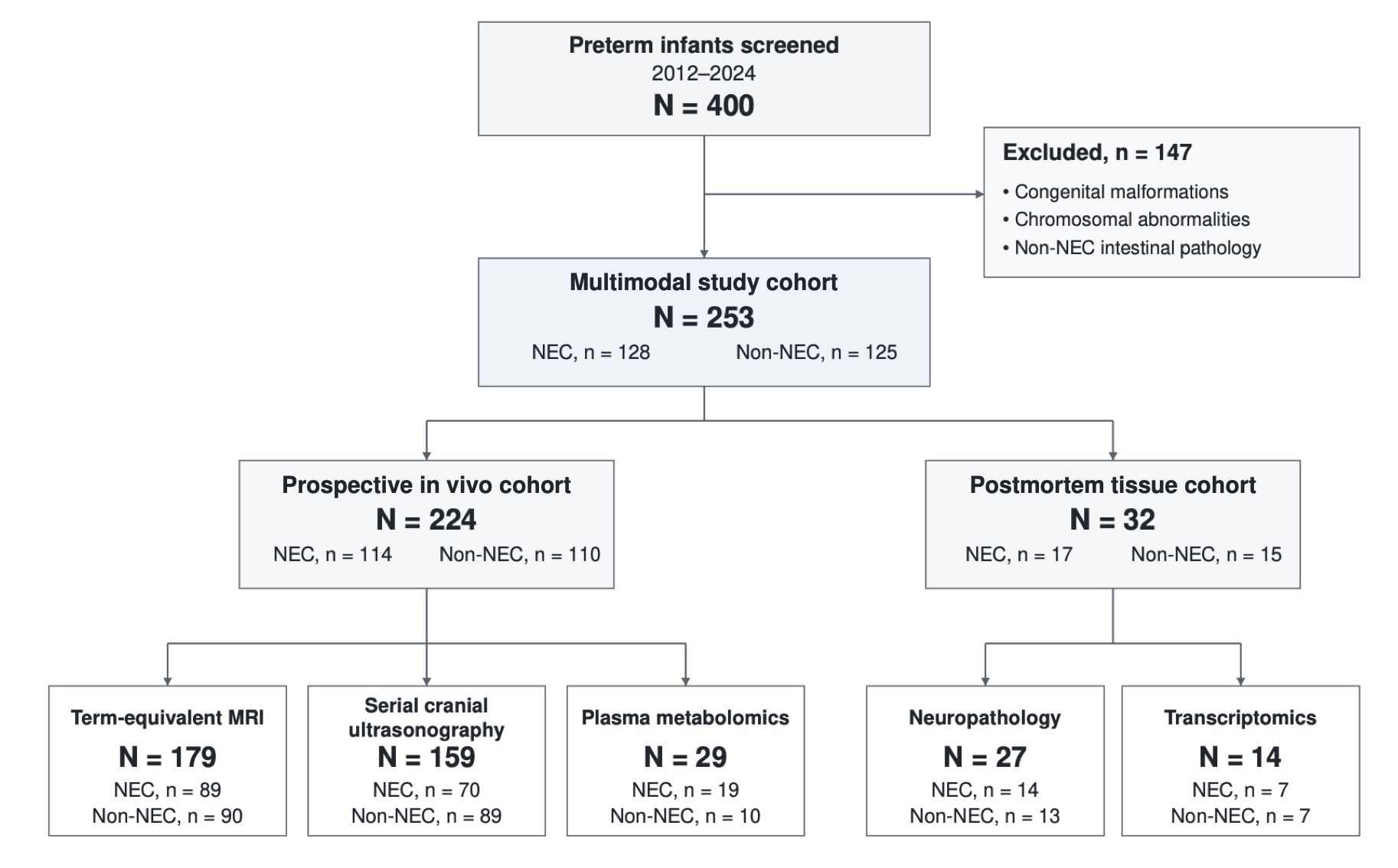

Flow diagram of participant screening and inclusion in the multimodal study. Of 400 preterm infants screened, 147 were excluded, yielding 253 unique infants (128 with NEC and 125 without NEC). The prospective in vivo cohort comprised 224 infants (114 with NEC and 110 without NEC), with overlapping subsets undergoing term-equivalent MRI (n = 179), serial cranial ultrasonography (n = 159), and plasma metabolomics (n = 29). The postmortem tissue cohort comprised 32 infants (17 with NEC and 15 without NEC), with overlapping subsets contributing to neuropathology (n = 27) and transcriptomics (n = 14), according to tissue availability. Three infants with NEC were included in both cohorts and were counted once in the overall study population. MRI indicates magnetic resonance imaging; NEC, necrotizing enterocolitis.

### eFigure 2. Lesion-Level Brain Abnormalities by Anatomical Structure

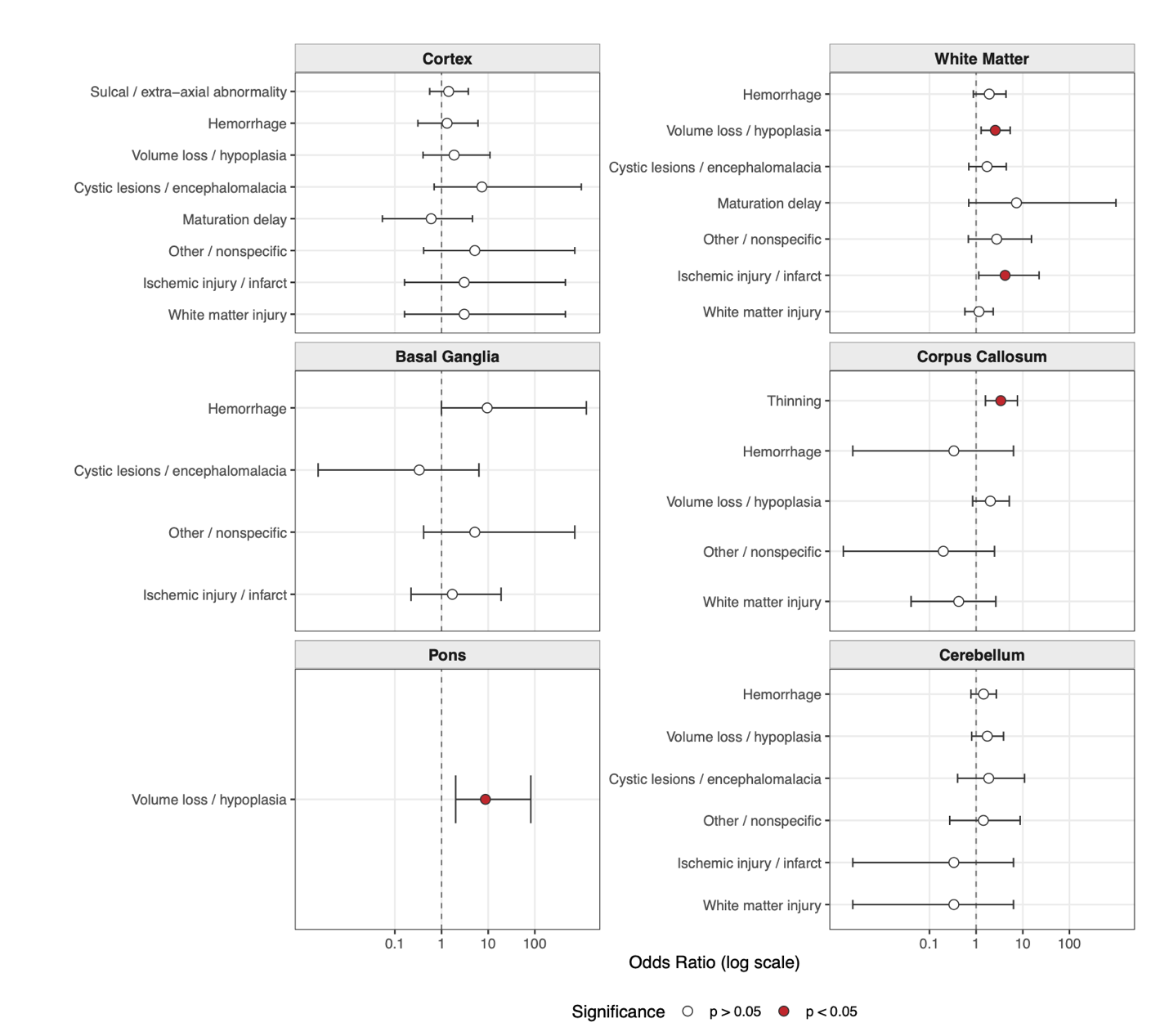

Forest plot showing odds ratios (ORs) for specific, lesion-level structural abnormalities on term-equivalent MRI in infants with NEC (n = 89) compared with non-NEC infants (n = 90). Structures are ordered rostral-to-caudal. Points represent ORs, and horizontal lines represent 95% CIs on a logarithmic scale. The vertical dashed line indicates an OR of 1.0. Red fill indicates a statistically significant association (*P* < .05), whereas white fill indicates a nonsignificant association (*P* ≥ .05). CI indicates confidence interval; MRI, magnetic resonance imaging; NEC, necrotizing enterocolitis; OR, odds ratio.

### eFigure 3. Regional Brain Abnormalities in Surgically vs Medically Managed Necrotizing Enterocolitis

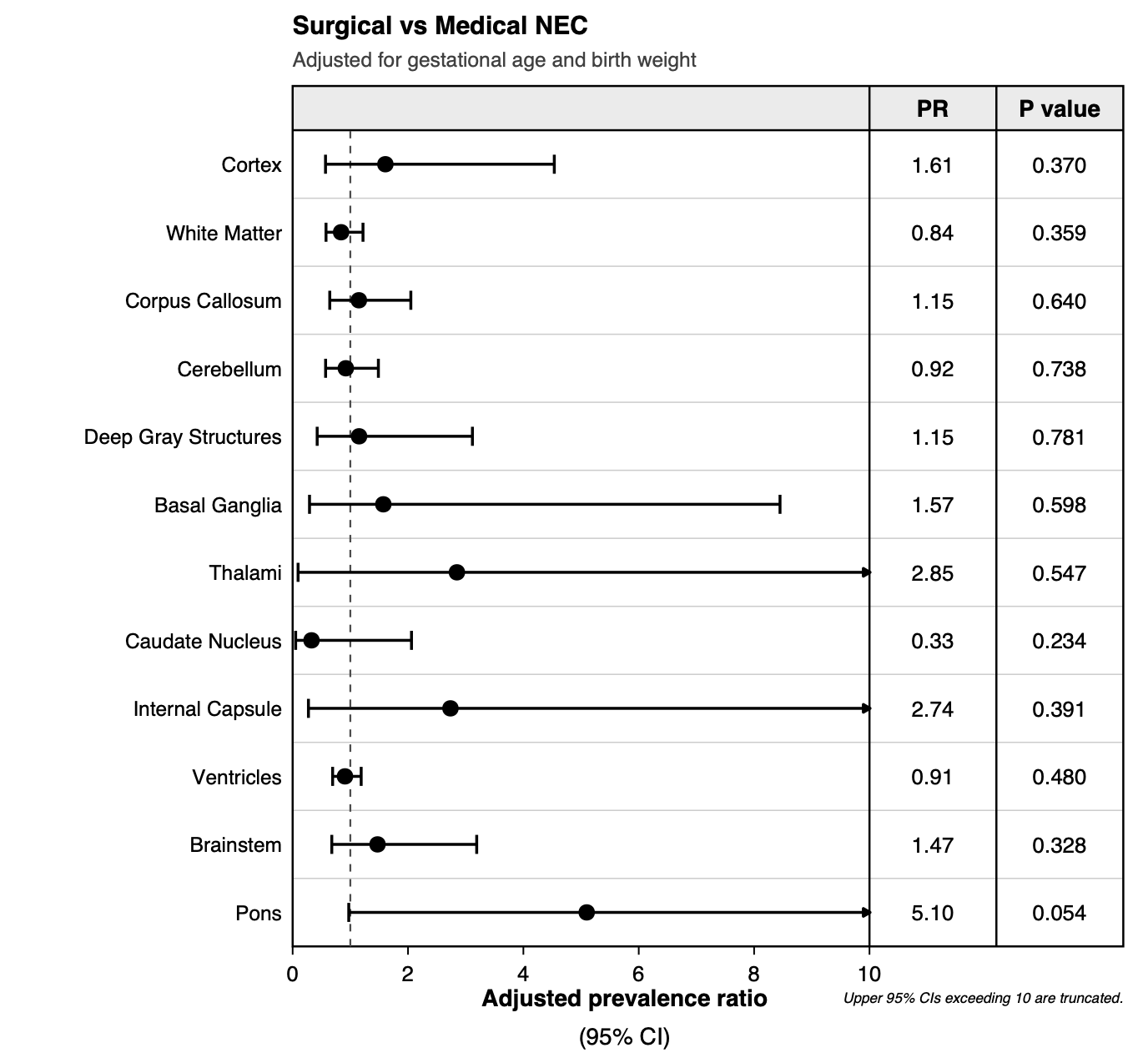

Forest plot displaying adjusted prevalence ratios (aPRs) and 95% CIs for regional brain abnormalities. The analysis compares infants with surgically managed NEC (n = 40) against those with medically managed NEC (n = 49). Models were adjusted for gestational age and birth weight. Point estimates (black circles) represent the aPR, with horizontal lines indicating the 95% CIs. Arrows pointing to the right indicate that the upper bound of the 95% CI exceeds 10 and has been truncated for visualization purposes. CI indicates confidence interval; NEC, necrotizing enterocolitis; aPR, adjusted prevalence ratio.

### eFigure 4. Intestinal Histopathology in the Postmortem Necrotizing Enterocolitis Cohort

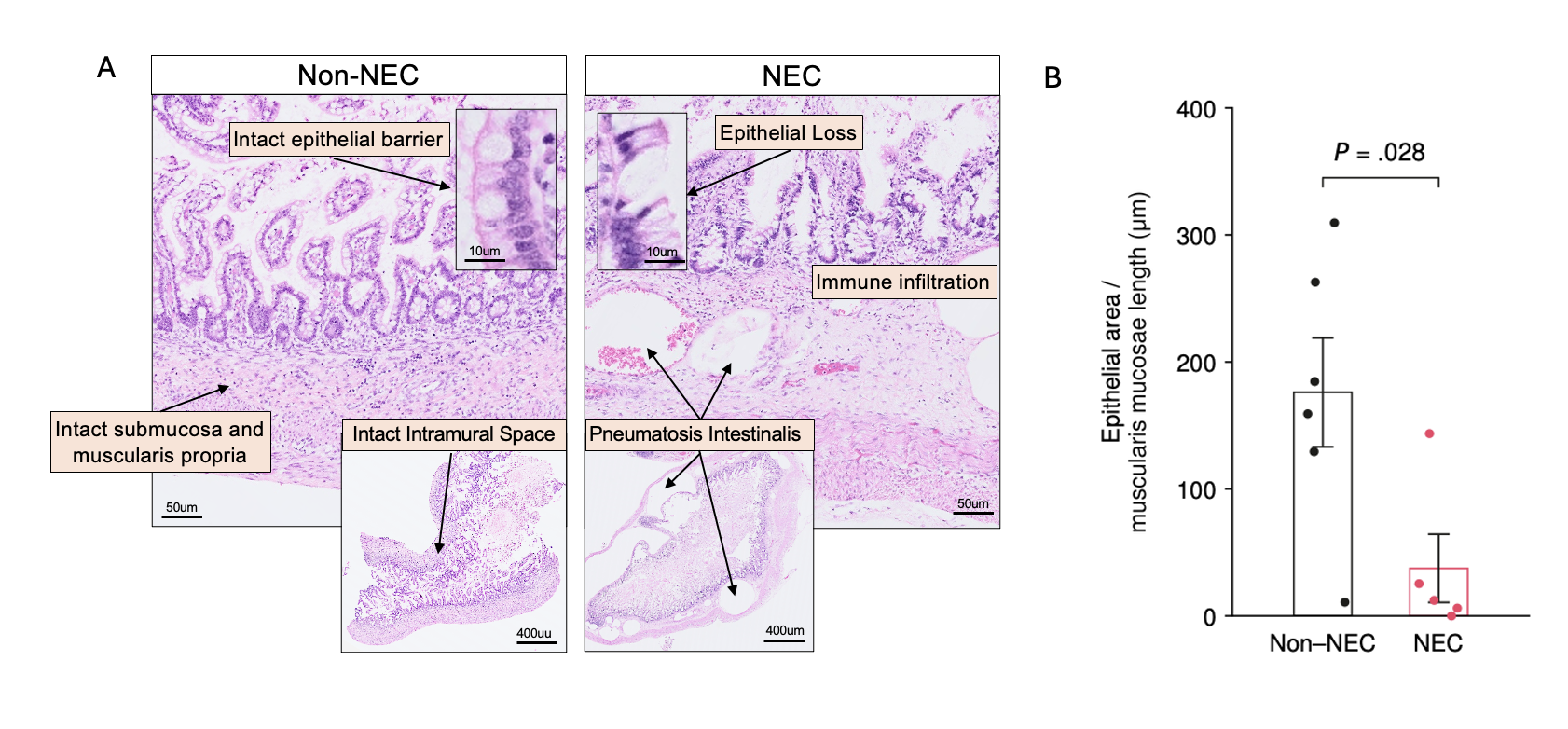

A, Representative hematoxylin-eosin–stained terminal ileum sections from infants without and with NEC. Non-NEC tissue demonstrates preserved epithelial and bowel wall architecture, whereas NEC tissue demonstrates epithelial loss, inflammatory infiltration, and pneumatosis intestinalis. Scale bars indicate 50 μm in the main images, 400 μm in the low-magnification overviews, and 10 μm in the high-magnification insets. B, Epithelial area normalized to muscularis mucosae length in NEC (n = 5) and non-NEC (n = 6) cases. Each point represents 1 infant, with epithelial areas and muscularis mucosae lengths pooled across available images before calculating the ratio. Bars indicate means, and error bars indicate SEMs. Black denotes non-NEC infants and red denotes infants with NEC. Groups were compared using a 2-sided exact studentized permutation test (*P* = .028). NEC indicates necrotizing enterocolitis.

### eFigure 5. Histopathological Assessment of Cerebral Subcortical White Matter

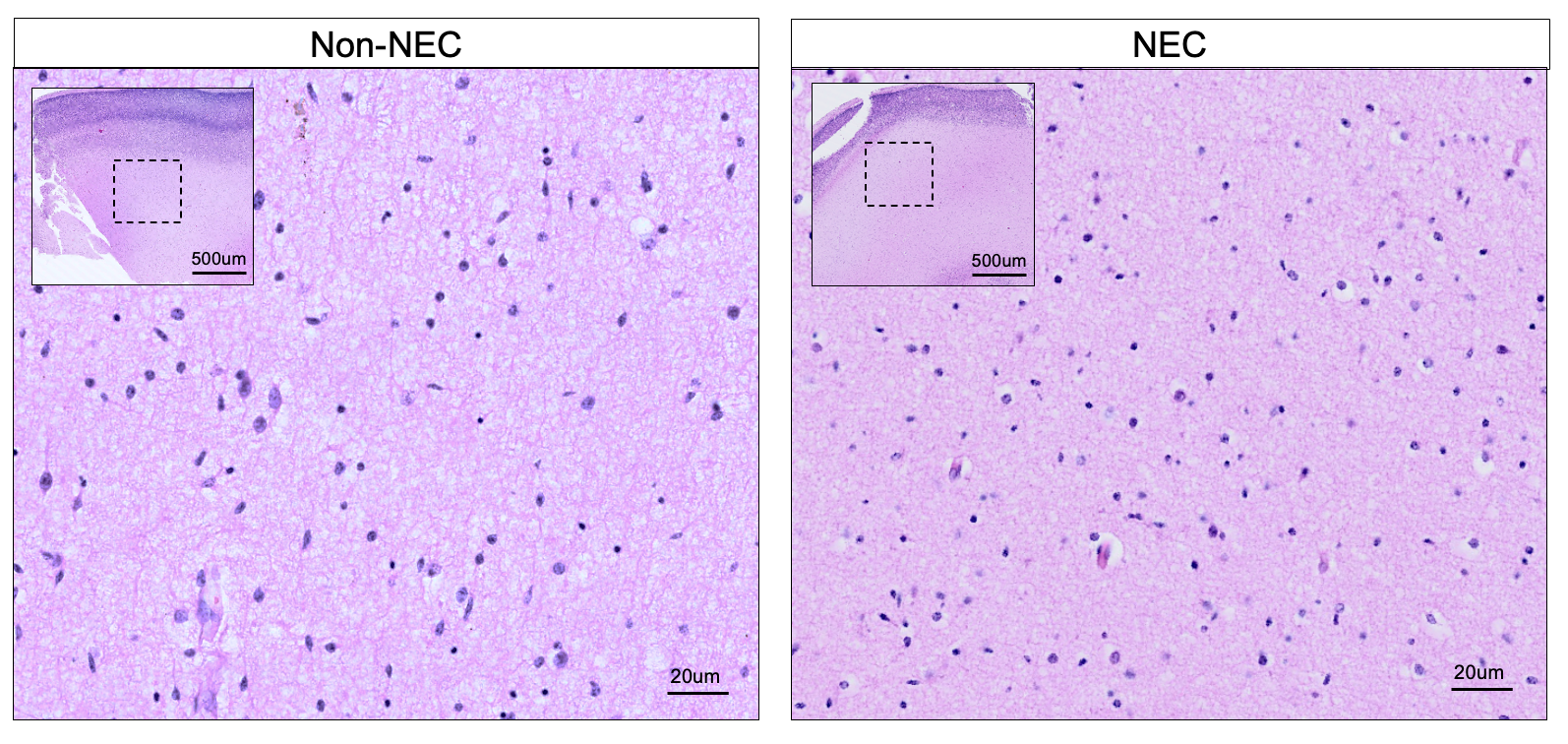

Representative hematoxylin-eosin–stained subcortical white matter sections from non-NEC (left) and NEC (right) preterm infants. Low-magnification insets delineate the sampled subcortical white matter (dashed boxes). High-magnification views demonstrate comparable prematurity-associated tissue rarefaction and axonal attenuation between groups. Scale bars indicate 500 μm in the insets and 20 μm in the main panels. NEC indicates necrotizing enterocolitis.

### eFigure 6. Cerebellar Cortical Injury and Morphometry in Necrotizing Enterocolitis

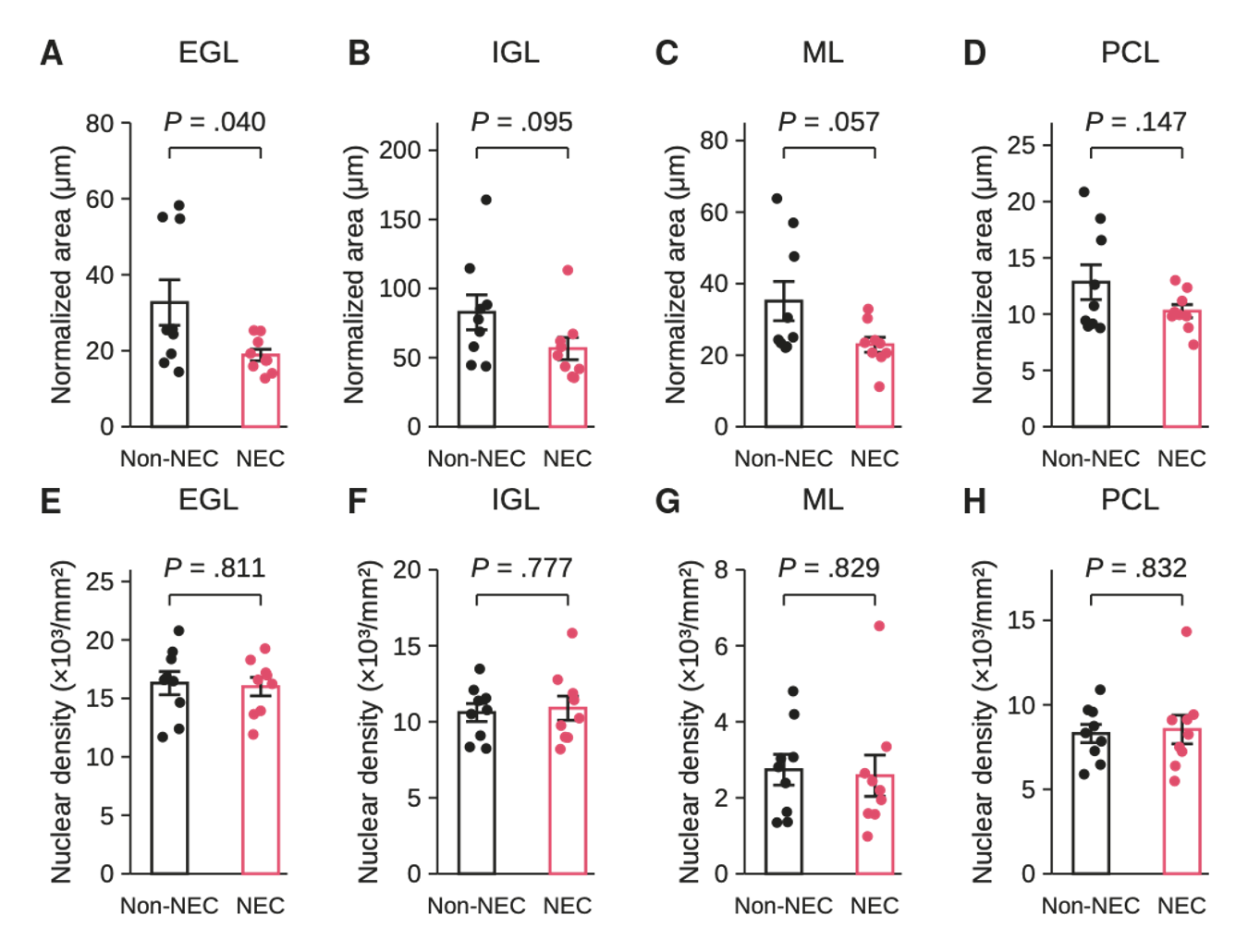

A–D, Normalized areas of the external granular layer (EGL; A), internal granular layer (IGL; B), molecular layer (ML; C), and Purkinje cell layer (PCL; D). Layer areas are expressed per unit reference length (μm; eMethods). E–H, Corresponding nuclear densities in the EGL (E), IGL (F), ML (G), and PCL (H), quantified using StarDist-based nuclear detection and expressed as ×10³ nuclei/mm². Each point represents an individual infant (9 with necrotizing enterocolitis [NEC] and 9 without NEC); black denotes non-NEC and pink denotes NEC. Bars indicate means, and error bars indicate SEMs. Between-group comparisons used 2-sided exact studentized permutation tests based on the Welch *t* statistic. Displayed *P* values are unadjusted.

### eFigure 7. Pontine Transcriptional Response in Necrotizing Enterocolitis

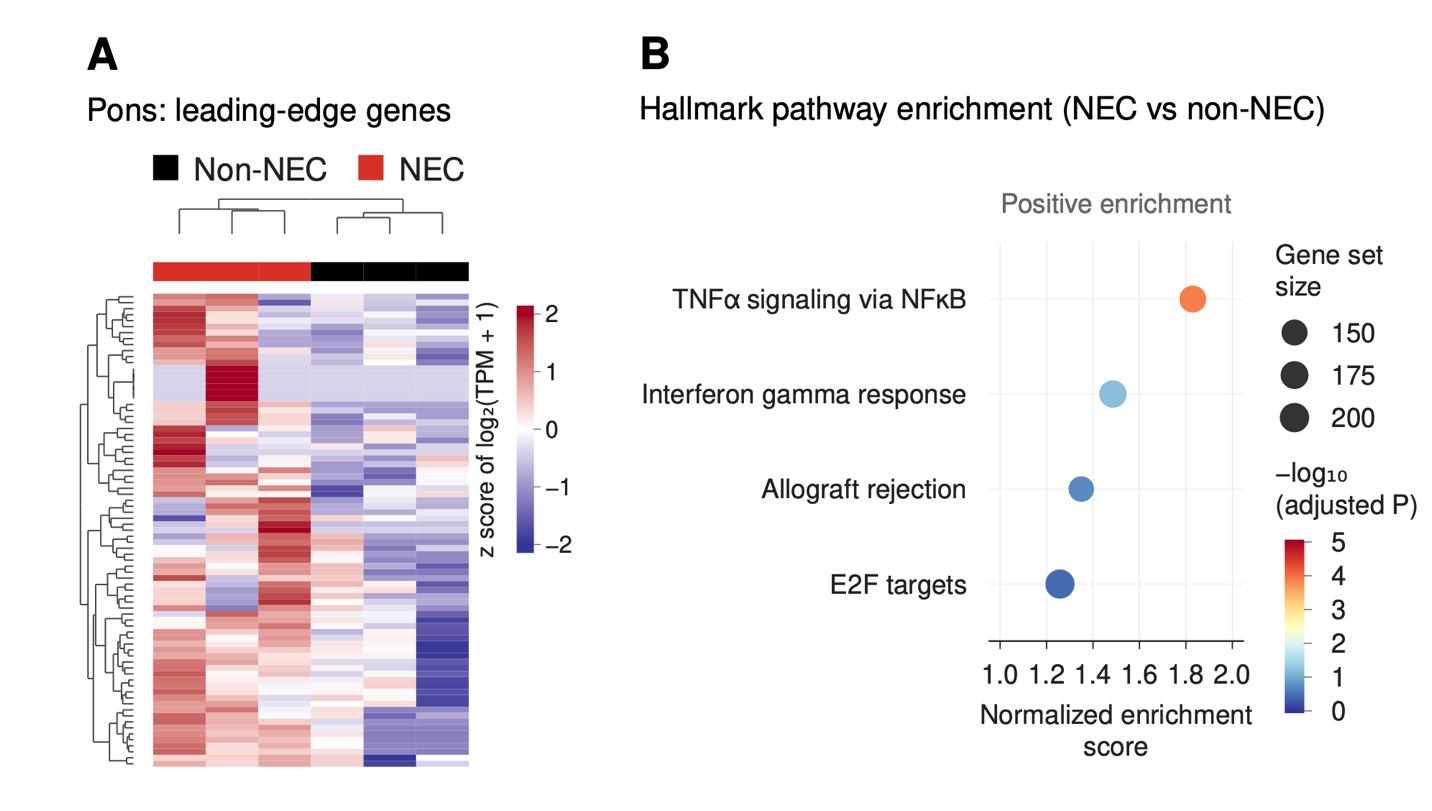

A, Hierarchically clustered heat map of pontine gene-expression patterns comparing NEC with non-NEC tissue. Expression values are displayed as gene-wise z scores of log₂-transformed transcript abundance, with clinical condition indicated above the heat map. B, Gene set enrichment analysis of selected Hallmark pathways with positive normalized enrichment scores in the NEC pons. Position on the x-axis represents the normalized enrichment score; circle size represents gene-set size, and color represents the negative log₁₀-transformed adjusted *P* value. NEC indicates necrotizing enterocolitis; NES, normalized enrichment score; NF-κB, nuclear factor kappaB; TNF, tumor necrosis factor.

### eFigure 8. Intestinal Transcriptional Programs in Necrotizing Enterocolitis

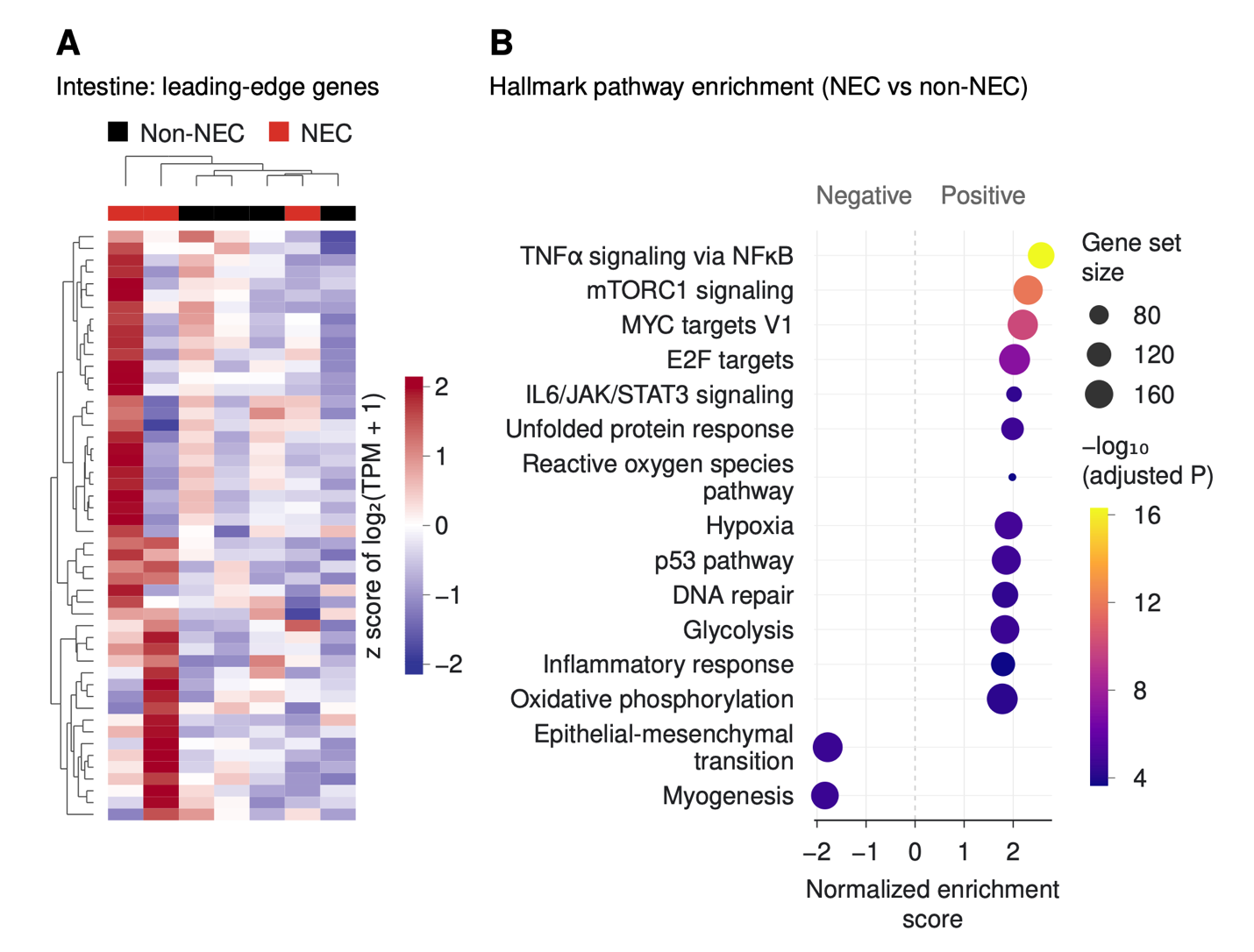

A, Heat map of leading-edge intestinal genes contributing to pathway enrichment in NEC compared with non-NEC tissue. Expression values are displayed as gene-wise z scores of log₂-transformed transcript abundance, with clinical condition indicated above the heat map. B, Hallmark pathway enrichment in NEC intestine. Position on the x-axis represents the normalized enrichment score; circle size represents gene-set size, and color represents the negative log₁₀-transformed adjusted *P* value. NEC indicates necrotizing enterocolitis; NES, normalized enrichment score; NF-κB, nuclear factor kappaB; TNF, tumor necrosis factor.

### eFigure 9. Candidate Intestine-to-Hindbrain Ligand-Receptor Signaling in Necrotizing Enterocolitis

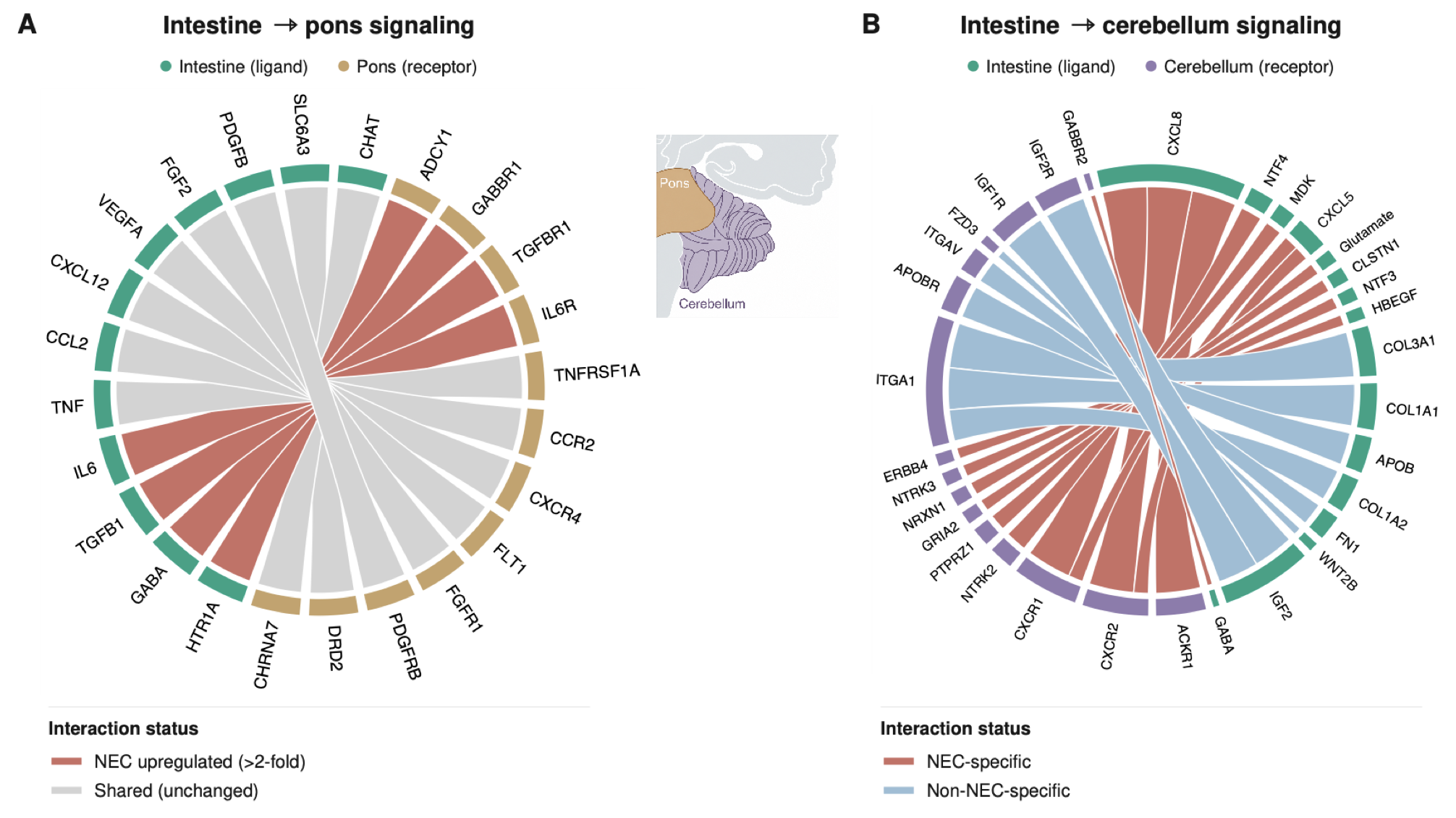

Cross-tissue ligand-receptor relationships were inferred from intestinal and regional brain transcriptomic profiles. A, Chord diagram showing candidate relationships between intestinal ligands and pontine receptors. Red connections indicate predicted interactions classified as increased in NEC (>2-fold; descriptive comparison without formal statistical testing), and gray connections indicate shared interactions classified as unchanged. B, Chord diagram showing candidate relationships between intestinal ligands and cerebellar receptors. Red connections indicate predicted interactions detected only in the NEC network, and blue connections indicate predicted interactions detected only in the non-NEC network. Green perimeter segments represent intestinal ligands; tan and purple segments represent pontine and cerebellar receptors, respectively. These expression-based predictions identify candidate relationships and do not establish functional intestine-to-brain signaling. NEC indicates necrotizing enterocolitis.

### eFigure 10. Cerebellar Transcriptional Suppression in Necrotizing Enterocolitis

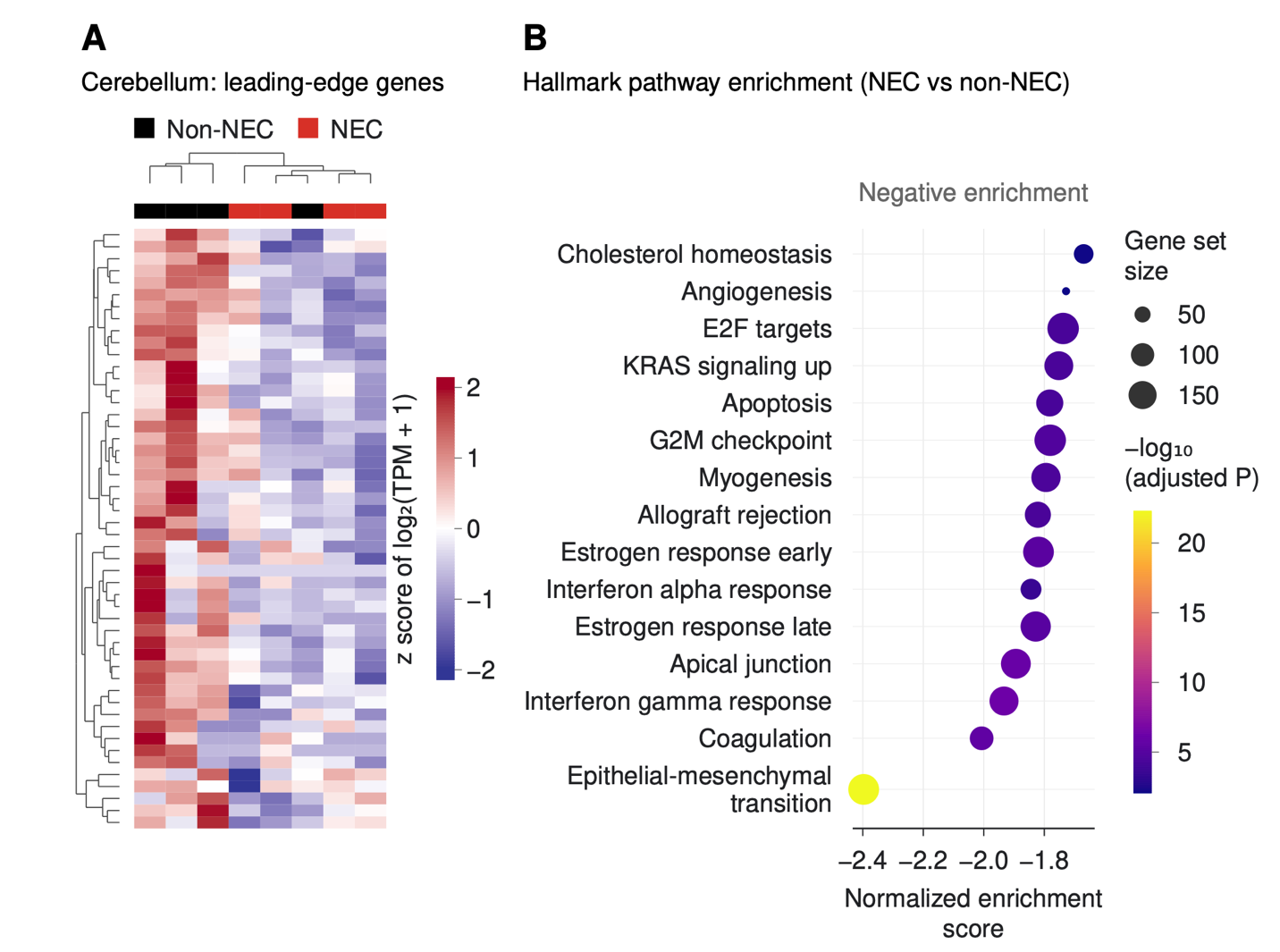

A, Heat map of leading-edge cerebellar genes contributing to pathway differences between NEC and non-NEC tissue. Expression values are displayed as gene-wise z scores of log₂-transformed transcript abundance, with clinical condition indicated above the heat map. B, Hallmark pathway enrichment analysis demonstrating predominantly negative enrichment in the NEC cerebellum. Position on the x-axis represents the normalized enrichment score; circle size represents gene-set size, and color represents the negative log₁₀-transformed adjusted *P* value. NEC indicates necrotizing enterocolitis; NES, normalized enrichment score.

### eFigure 11. Global Cranial Trajectories and Growth Velocities in Necrotizing Enterocolitis

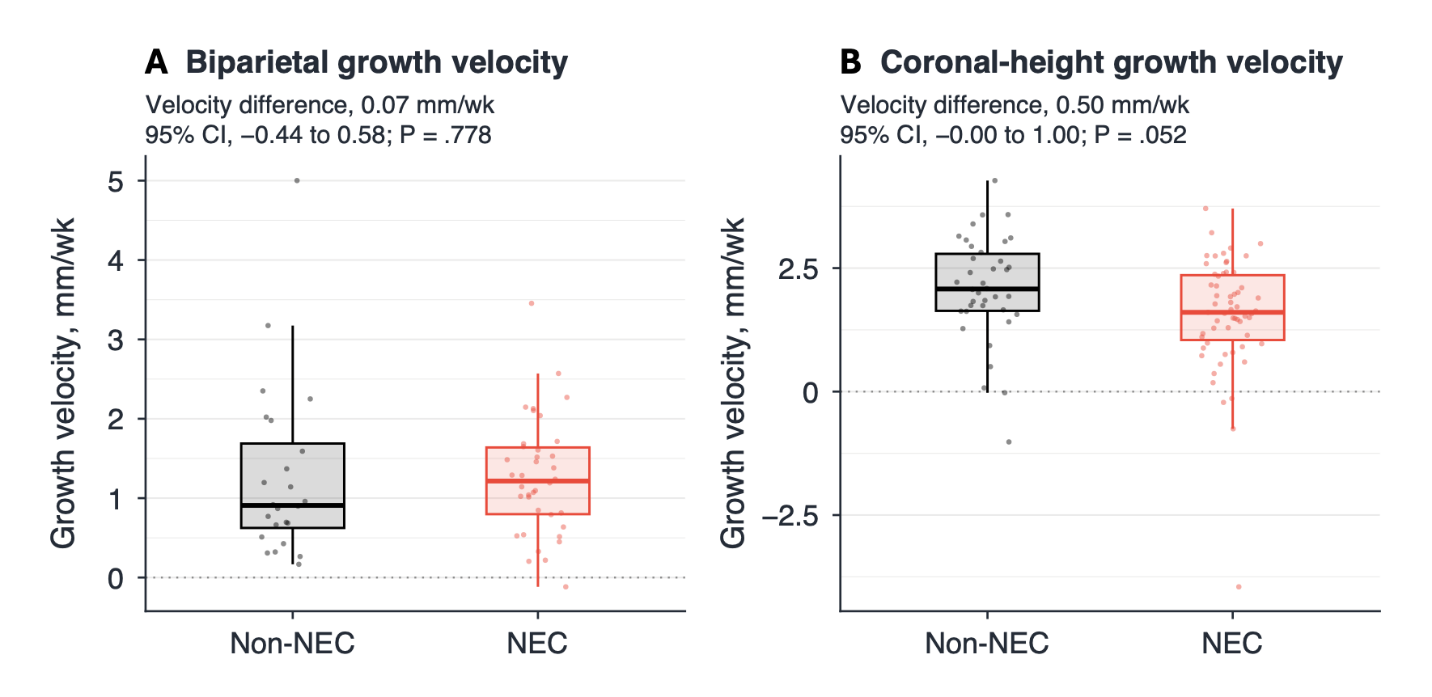

A and B, Box plots comparing biparietal diameter (A) and brain coronal-height (B) growth velocities between infants without and with NEC. Analyses included 62 infants for biparietal diameter (38 with NEC and 24 without NEC) and 97 infants for coronal height (59 with NEC and 38 without NEC). Black denotes non-NEC infants and red denotes infants with NEC. Points show observed growth velocities; boxes represent the interquartile range, with the horizontal line indicating the median. Whiskers extend to the most extreme value within 1.5 times the interquartile range below the first quartile or above the third quartile. Between-group differences were estimated using analysis of covariance adjusted for midpoint postmenstrual age and gestational age at birth. The adjusted difference in growth velocity, expressed as non-NEC minus NEC, was 0.07 mm/week for biparietal diameter (95% CI, −0.44 to 0.58; *P* = .778) and 0.50 mm/week for coronal height (95% CI, −0.004 to 1.005; *P* = .052). CI indicates confidence interval; NEC, necrotizing enterocolitis.
